# Isthmus progenitor cells contribute to homeostatic cellular turnover and support regeneration following intestinal injury

**DOI:** 10.1101/2022.04.26.489611

**Authors:** E. Malagola, A. Vasciaveo, Y. Ochiai, W. Kim, M. Middelhoff, H. Nienhüser, L. Deng, B. Belin, J. LaBella, LB. Zamechek, M.H. Wong, L. Li, C. Guha, CW. Cheng, KS. Yan, A. Califano, T.C. Wang

**Affiliations:** Division of Digestive and Liver Diseases, Department of Medicine and Irving Cancer Research Center, Columbia University Medical Center, New York, NY 10032, USA; Department of Systems Biology, Columbia University, New York, NY 10032, USA; Klinik und Poliklinik für Innere Medizin II, Klinikum rechts der Isar, Technische Universität München, Munich, Germany; Department of General, Visceral and Transplant Surgery University Hospital Heidelberg Im Neuenheimer Feld 420 69120, Heidelberg, Germany; Stowers Institute for Medical Research, Kansas City, MO 64110, USA; Department of Pathology and Laboratory Medicine, University of Kansas Medical Center, Kansas City, KS 66107, USA; Department of Cell, Developmental & Cancer Biology, Oregon Health & Sciences University 3181 SW Sam Jackson Park Road, L215, Portland, USA; Department of Radiation Oncology, Albert Einstein Cancer Center, Albert Einstein College of Medicine,1300 Morris Park Avenue, Bronx, NY 10461, USA; Columbia Stem Cell Initiative, Department of Genetics and Development, Columbia University Irving Medical Center, New York, New York; Department of Medicine, Vagelos College of Physicians and Surgeons, Columbia University Irving Medical Center, New York, NY 10032, USA; Department of Biochemistry & Molecular Biophysics, Vagelos College of Physicians and Surgeons, Columbia University Irving Medical Center, New York, NY 10032, USA; Department of Biomedical Informatics, Vagelos College of Physicians and Surgeons, Columbia University Irving Medical Center, New York, NY 10032, USA; Herbert Irving Comprehensive Cancer Center, Columbia University Irving Medical Center, New York, NY 10032, USA

## Abstract

The currently accepted intestinal epithelial cell organization model proposes that crypt base columnar (CBC) cells marked by high levels of *Lgr5* expression represent the sole intestinal stem cell (ISC) compartment. However, previous studies have indicated that Lgr5^+^ cells are dispensable for intestinal regeneration, leading to two major hypotheses: one favoring the presence of a quiescent reserve stem cell population, the other calling for differentiated cell plasticity. To investigate these possibilities, we studied crypt epithelial cell organization, during homeostasis and regeneration, in unbiased fashion, via high-resolution single-cell profiling. These studies, combined with *in vivo* lineage tracing, show that *Lgr5* is not a specific ISC marker and that stemness potential exists beyond the crypt base in the isthmus region, whose cells, contrary to differentiated cells, participate in tissue homeostasis and support intestinal regeneration. Our results provide a novel model of organization for the intestinal crypt epithelium in which stemness potential is not restricted to CBC cells and suggesting that neither de-differentiation nor reserve stem cell populations are drivers of intestinal regeneration.

## Introduction

The intestinal epithelium is characterized by a high cellular turnover rate, making it an attractive model to study adult stem cell biology. The intestinal stem cell (ISC) is defined by its ability to self-renew and to give rise to all intestinal epithelial lineages. Lineage tracing studies have uncovered a multitude of genes expressed in ISCs^1–7^; among others, *Lgr5* has been widely accepted as the primary ISC marker^8^ due to its proposed high degree of specificity. In particular, *Lgr5* was reported to be selectively expressed in crypt base columnar (CBC) cells, located at the very bottom of the intestinal crypts, between Paneth cells, typically at positions +1 to +3^8^. However, recent studies on intestinal regeneration following high doses of radiation^5, 6, 8^, or selective *Lgr5*-expressing cell ablation^9^, have challenged this view by showing that other crypt epithelial cells can compensate for the loss of Lgr5^+^ cells. These observations suggest that the ‘Lgr5-CBC’ model may not fully represent the regenerative ability of the intestinal epithelium. To account for these findings, two major hypotheses have emerged: the first proposing that, in addition to the Lgr5^+^ ISCs, a quiescent ‘+4’ stem cell population may act as a ‘reserve’ to replenish the ISC pool following intestinal injury^5, 7, 9^, the second asserting that, following ISC loss, multiple differentiated cell types may undergo de-differentiation and serve as *bona fide* ISCs^10–13^.

Single cell transcriptomics has emerged as a powerful tool to study tissue heterogeneity and to uncover the identity and dynamics of rare cell populations^14, 15^. With respect to the intestine, this approach has been instrumental in detailing the specific differentiation states of diverse intestinal lineages, as well as to investigate the cellular composition of the regenerating intestine^7, 16–18^. However, previous studies and descriptions of the intestinal epithelium hierarchical organization have relied heavily on established markers, thus forcing an interpretation compatible with prior ISC models and limiting the generation of alternative hypotheses^19^.

Recently, a number of studies have shown that computational cell potency inference is effective in reconstructing cell hierarchies within tissues^20–23^. These novel approaches can infer the hierarchical organization of intestinal epithelial cells *de novo*, in unbiased fashion, *i.e.*, without having to rely on prior knowledge. Such an approach, coupled with relevant experimental assays, can thus help assess whether the current ISC models are consistent with actual tissue organization. Furthermore, these methodologies allow the investigation of putative changes in intestinal epithelial cell potency following intestinal injury, as well as the possible emergence of a reserve stem cell population.

To address these questions, we studied intestinal crypt epithelial cells by combining single cell profiling with novel lineage tracing mouse and then analyzed their organization using an unbiased, data-driven approach that does not rely on previously established markers. Our data show that *Lgr5* overlaps with but is not uniquely expressed in stem and progenitor cells. More critically, they indicate that stemness potential exists beyond the CBC cells’ compartment in the isthmus region. Based on these results, we propose a novel model of intestinal organization in which additional actively cycling cells have the potential to act as the multipotent, self-renewing ISCs and can contribute to homeostatic cellular turnover alongside CBC cells. Following intestinal injury, surviving isthmus cells guide intestinal regeneration, identifying them as the postulated reserve ISCs, and suggesting that de-differentiation constitutes an exceptional event rather than the main driver of intestinal regeneration.

## Results

### An unbiased approach to elucidate the organization of intestinal crypt epithelial cells

To study intestinal crypt epithelial cell composition, we developed an integrative, experimental and computational pipeline designed to recover the transcriptional identity of individual intestinal cells, as well as their hierarchical relationships by transcriptomic and epigenetic single cell profile analysis (Fig. 1a). A high level of enrichment of crypt epithelial cells was achieved via nearly complete villus epithelial cell exclusion, using a villi-cell specific antibody (B6A6^24^) (Extended Data Fig. 1a).

**Figure 1.**
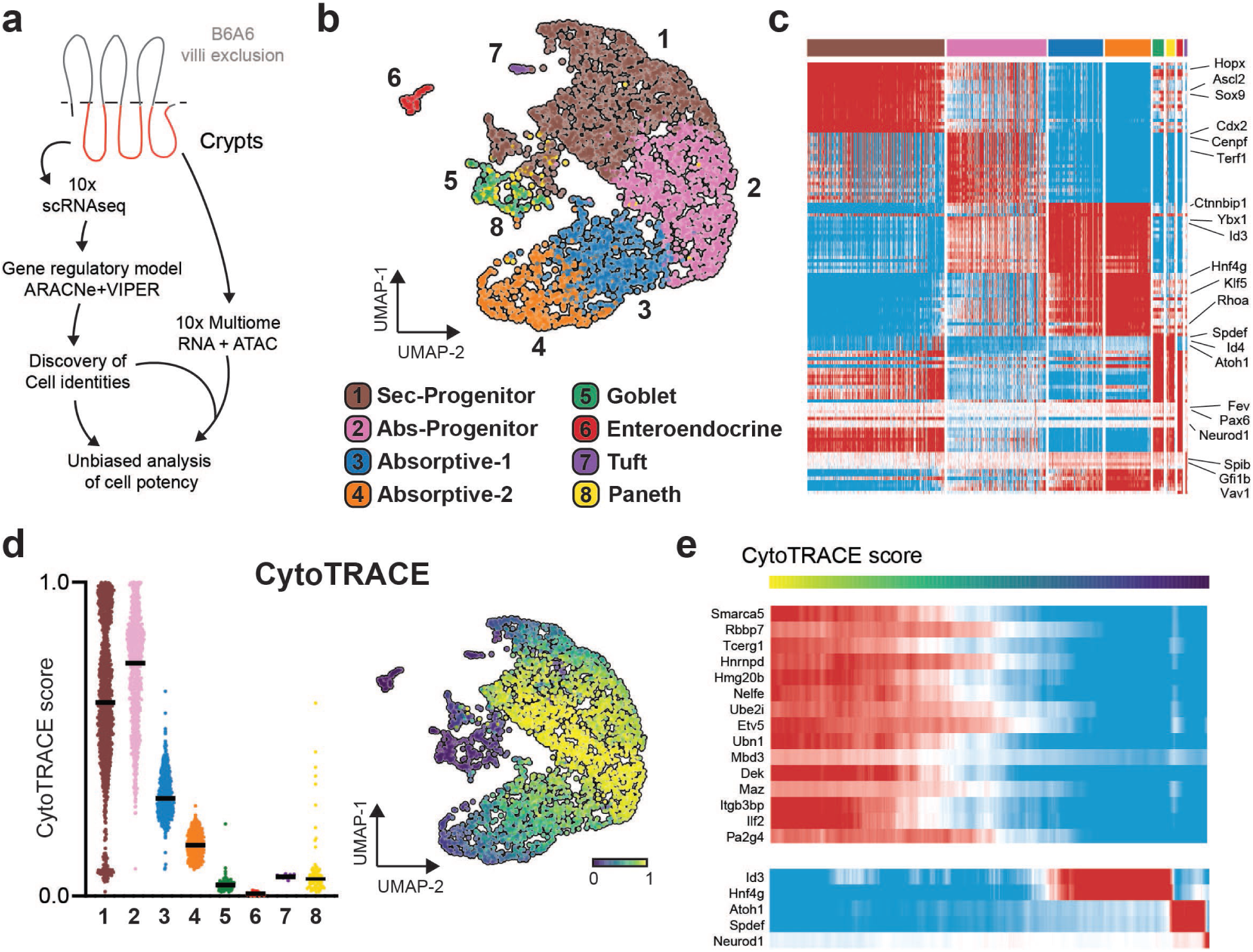
An unbiased approach to elucidate the organization of intestinal crypt epithelial cells: a. Schematic representation of the experimental workflow and computational pipeline for the analysis of crypt epithelial cells. (*See methods*) b. UMAP plot showing protein activity-based clustering solution based on gene regulatory network analysis. c. Heatmap showing top differentially activated regulatory proteins. (Active scores in red) d. Violin- and UMAP plots showing CytoTRACE scores for individual cells; black lines indicate median value per cluster. e. Pseudotime analysis using CytoTRACE scores to order cells based on inferred cell potency (high – yellow - to low – blue - potency from left to right), showing protein activity changes of top correlating regulatory proteins. On the bottom, protein activity profile for known transcription factors involved in cellular differentiation. (Active scores in red)

To avoid critical gene dropout issues common to single cell RNA-seq (scRNA-seq) profile analyses—where, on average, ≥ 80% of the genes are undetectable in each individual cell^25^—we relied on VIPER, an established network-based algorithm designed to measure the transcriptional activity of regulatory and signaling proteins from scRNA-seq profiles^26^. Specifically, akin to highly multiplexed gene reporter assay, VIPER measures a protein’s activity based on the differential expression of its 50 highest-confidence transcriptional targets (*regulon*), as identified by the ARACNe algorithm^27, 28^. Critically, we have shown that this methodology allows identification of subpopulations that are essentially undetectable by gene expression analysis^14, 15^, as well as robust, highly reproducible quantification of protein activity, which compares favorably with antibody-based single cell measurements^14^, as well as removal of technical artifacts and batch effects^29^.

To identify intestinal stem and progenitor cells, we combined analyses of crypt epithelial cell chromatin accessibility via single-cell Assay for Transposase-Accessible Chromatin scATAC-seq^30^ and cell potency inference via scRNA-seq analysis^20, 22, 23^ (*See Methods*). Three major subpopulations emerged initially from cluster analyses of >3,500 crypt epithelial cells, corresponding to two established intestinal lineages, secretory (*Hepacam2^hi^*, *Tff3^hi^*, etc.) and absorptive (*Ckmt1^hi^*, *Ccl25^hi^,* etc.) cells, with a third subpopulation mostly comprised of stem and progenitor cells (*Stmn1^hi^*, *Dek^hi^*, etc.)^16, 18, 19^ (Extended Data Fig. 1b-f, Extended Data Table 3-4). Importantly, inference of cell potency, using independent algorithms^20, 21, 23^, showed a high degree of similarity and correlated well with the degree of chromatin accessibility observed, supporting the effectiveness of these methodologies.

To further reveal the lineage-specific nature of intestinal sub-populations, we performed independent sub-clustering of these three major populations (Fig. 1b-c, Extended Data Fig. 1g, Extended Data Table 5-6). This refined analysis revealed two absorptive clusters, corresponding to subpopulations representing early specified and more committed cells, respectively, as well as three clusters corresponding to established secretory subpopulations, including tuft (*Dclk1*+), enteroendocrine (*NeuroD1*+), and goblet (*Atoh1*+) cells^19^. Notably, consistent with previous single cell, VIPER-based, studies^14, 15^, the vast majority of subpopulation-specific proteins—including established intestinal lineage markers such as Fev^31^, and Gfi1b^19^—could only be detected by VIPER-based protein activity analysis but not based on the differential expression of their encoding genes (Extended Data Fig. 1h). Due to their relative small number, *Lyz1*^hi^ Paneth cells did not form an independent cluster but rather were intermixed within the goblet cluster (Extended Data Fig. 2a-b). To overcome this limitation, we further identified Paneth cells based on expression of their established markers^16^ thus allowing for their inclusion in further analyses. Atoh1 and Spdef, both previously established regulators of Paneth cell’s identity, were among the most activated proteins in these cells, explaining their initial co-segregation within the goblet cells cluster.

**Figure 2.**
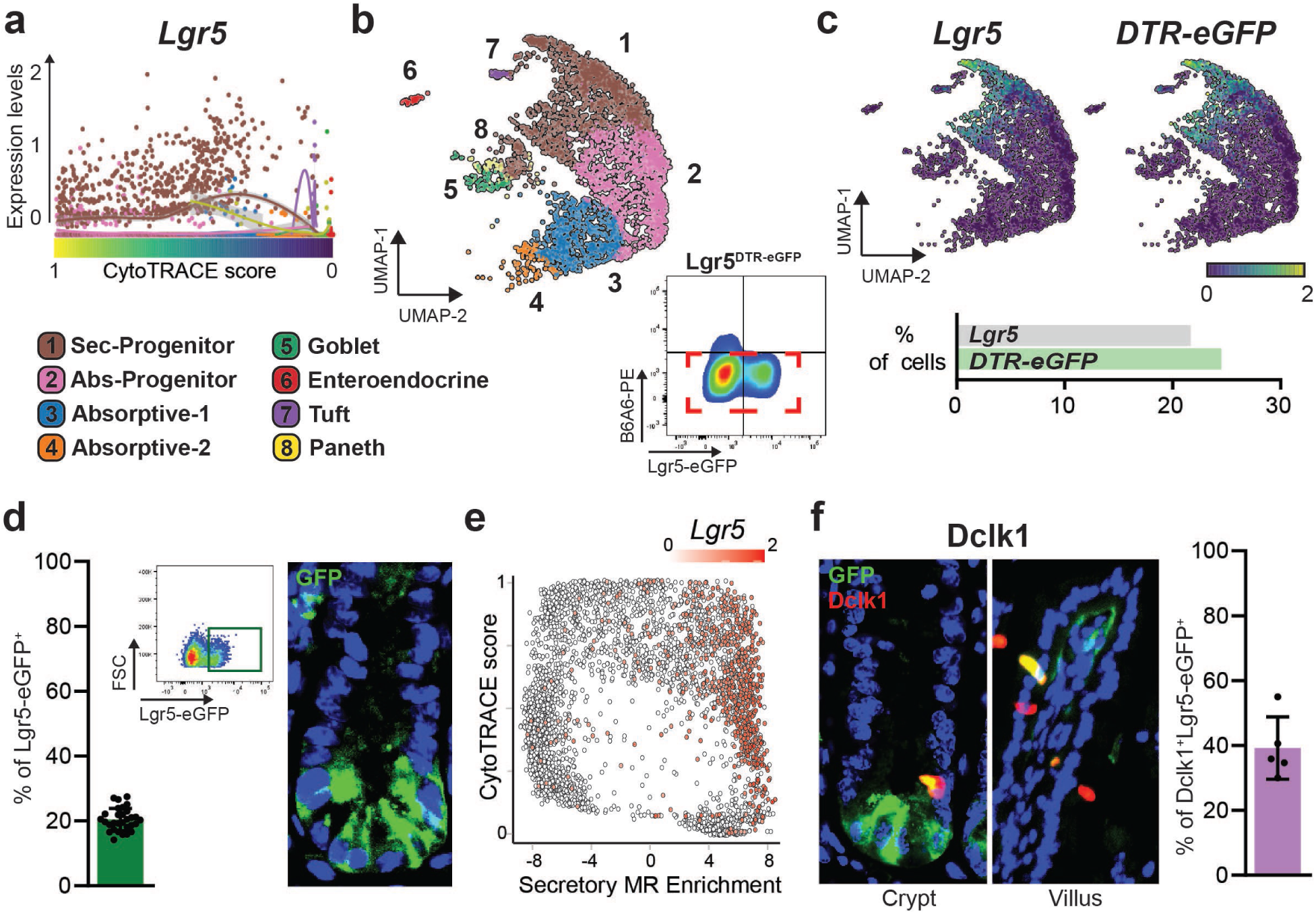
Lgr5 expression is not restricted to intestinal stem and progenitor cells: a. Pseudotime analysis using CytoTRACE scores to order cells based on inferred cell potency (high to low potency from left to right), showing expression levels of *Lgr5* on y-axis. Cells are colored based on their cluster. **b.** Sorting strategy and UMAP plot of Lgr5^DTR-eGFP^ crypt epithelial cells. (Sorted gate highlighted in Red) **c.** UMAP plots showing expression of *Lgr5*^Wt^ and *Lgr5^DTR-eGFP^* alleles, below bar-plot showing the number of cells expressing each allele in the dataset (as % of total cells). **d.** Bar-plot and gating strategy for flow cytometry –based quantification of Lgr5-eGFP^+^ cells in purified crypts (n=30); together with representative image of immunofluorescence staining for Lgr5-eGFP in mouse jejunum. **e.** Scatter plot of inferred cell potency (CytoTRACE) on y-axis and enrichment scores for Secretory lineage on x-axis. Cells are colored based on Lgr5 expression. **f.** Immunofluorescence staining for Dclk1 and bar-plot showing quantification of double positive cells (Dclk1^+^Lgr5-eGFP^+^ over total Dclk1^+^) (n=5).

Finally, stem and progenitor cells co-segregated within two clusters corresponding to subpopulations presenting similar levels of inferred potency, suggesting that this subdivision does not reflect a hierarchical partition of these cells (Fig. 1d, Extended Data Fig. 2c-e). Rather, the highest level of stemness potential was detected at the boundary between these two clusters (Extended Data Fig. 2f), suggesting that these reflect a gradual yet progressive differentiation of progenitor cells towards a secretory (Sec-Progenitor) or absorptive (Abs-Progenitor) fate, respectively. Consistent with these findings, discrimination of candidate ISCs from proliferative progenitors, by joint CytoTRACE and RNA content analysis, as previously described^20^, indicated that these candidate were well included in the top decile of inferred cell potency corresponding to virtually all boundary cells (Extended Data Fig. 2g). Analogous results were obtained when analyzing previously published datasets of crypt epithelial cells^7, 17^, which however presented a lesser degree of enrichment for stem and progenitor cells, corroborating the advantage of our isolation strategy (Extended Data Fig. 2h-i). Furthermore, lineage-specific enrichment analyses of crypt epithelial cells further highlighted the observed bias of the progenitor populations towards either a secretory or absorptive fate, respectively (Extended Data Fig. 2j).

Notably, these analyses provide a unique opportunity to study the molecular features that best describe stemness within the intestine. To this aim, we characterized the regulatory factors with the most statistically significant association with inferred cell potency, by correlating the protein activity profiles, as assessed by VIPER, with inferred potency, as assessed by CytoTRACE^20^, on a cell by cell basis (Fig. 1e, Extended Data Fig. 3, Extended Data Table 7). Regulatory proteins, whose activity presented the most significant correlation with CytoTRACE scores, were highly enriched for factors known to regulate stemness across different tissues, including the intestine^32–38^. Consistently, known regulators of intestinal differentiation^19^ had negative correlation scores.

**Figure 3:**
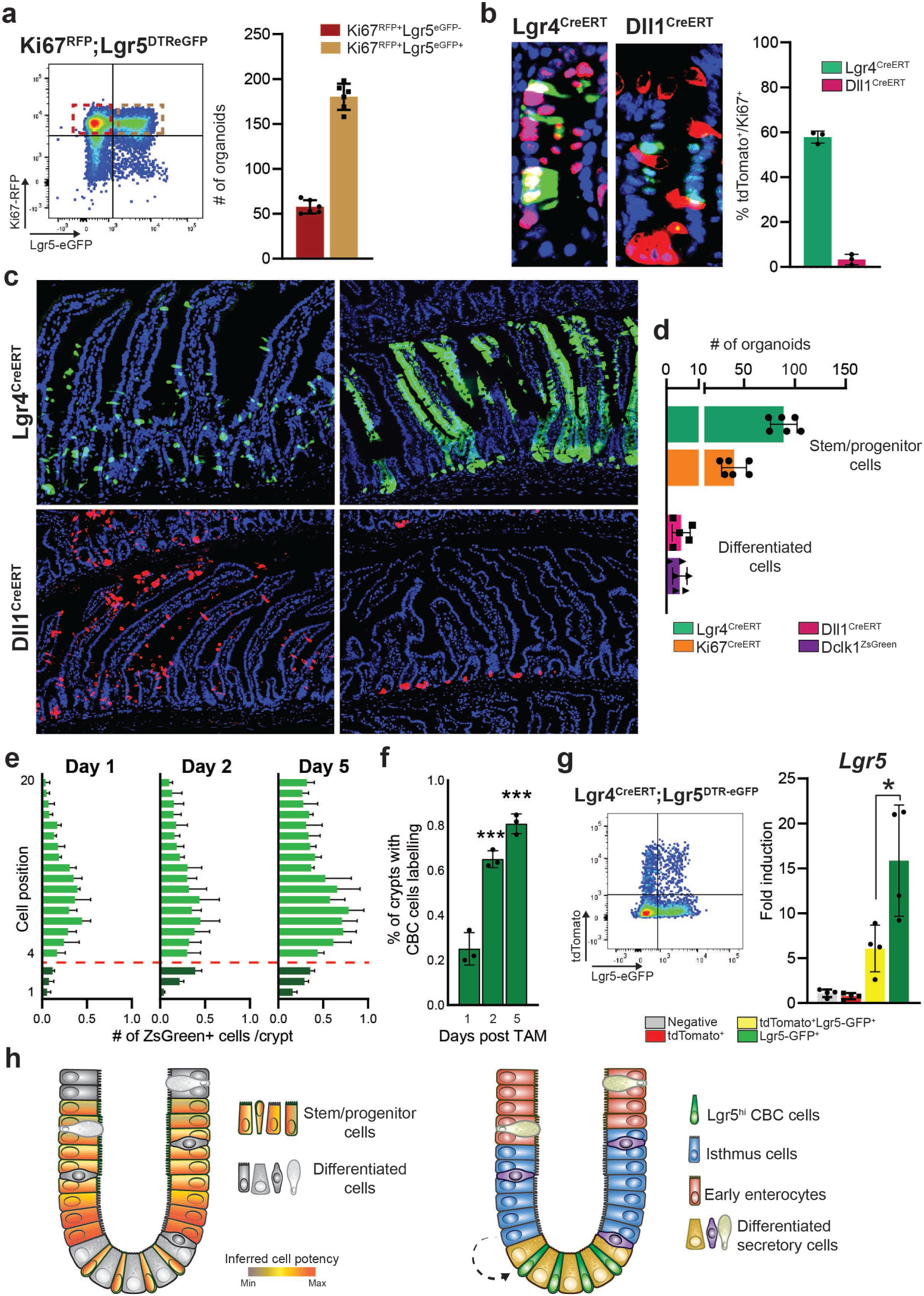
Isthmal stem cells sustain intestinal tissue homeostasis: a. Representative scatter-plot for flow cytometry –based cells sorting (Ki67^RFP^;Lgr5^DTReGFP^). On the right, bar-plot showing number of organoids generated from the isolated fractions. **b.** Representative image of Ki67 staining combined with lineage tracing in Lgr4^CreERT^ and Dll1^CreERT^ mice (24h post TAM). On the right, bar-plot showing quantification of % of Ki67+/tdTomato+ cells. **c.** Representative images of Lgr4^CreERT^ and Dll1^CreERT^ –derived labelling 1 day and 1 (Dll1^CreERT^) or 6 (Lgr4^CreERT^) months post TAM induction. **d.** Bar-plot showing quantification of the number of organoids generated from each sorted population. **e.** Bar-plots showing quantification of traced cells in Lgr4^CreERT^ based on cell position within the crypt. Red dotted line used to mark the boundary between CBC and isthmus cells (n=3). **f.** Bar-plots showing percentage of crypt-units with at least one labelled CBC cell in Lgr4^CreERT^. **g.** Flow cytometry analysis of overlap between Lgr4^CreERT^ labelled cells (24h post TAM) and Lgr5^eGFP+^ cells; together with bar plot showing expression levels of *Lgr5* in single sorted cells, groups labelled in the panel. (n=4). **h.** (Left) Schematic representation of the intestinal crypts, cells are colored based on inferred cell potency. Highest stemness potential is found in the isthmus region (e.g. Lgr5^low^ cells). (Right) Schematic representation of intestinal crypts, isthmus cells, colored in blue, have the capacity to migrate downward and give rise to Lgr5+ CBCs during homeostasis. Statistical method: Unpaired t-test, two-tailed; *: p<0.05, **: p<0.01, ***: p<0.01, ****: p<0.01.

Among others, the transcription factor Yy1, was previously reported to play an essential role in preventing ISC differentiation and to impair epithelial cells organoids forming capacities upon genetic knock-out^36^. Several chromatin remodeler enzymes also emerged from the analysis— including Smarca5^33^ and Atad2^34^, which have been previously associated with stemness—thus highlighting the crucial role of chromatin remodeling in cell potency. Several splicing factors also emerged within the broad group of transcriptional regulators associated with cell potency, including members of the Srsf family, which were recently shown to play an important role in regulating the intestinal epithelium^37^.

Focusing on genes whose differential expression, rather than activity, correlated with inferred cell potency (Extended Data Table 8), we identified *Smarca5*, *Stmn1* and *Dek*, all representing previously proposed markers of adult stem cells^32, 39^. Birc5a^40^ and Ncl^41^, both described as highly expressed in embryonic stem cells, also emerged among the most correlated genes. Finally, *Fgfbp1* was also positively correlated with inferred cell potency (*Capdevila C et al., accompanying paper*). These analyses provide an unbiased characterization of intestinal crypt epithelial cells using mechanism-based dissection of protein activity in single-cell data, via regulatory network analysis, and highlight novel factors that best recapitulate intestinal cell potency.

### Lgr5 expression is not restricted to intestinal stem and progenitor cells

Lgr5 has been described as a highly specific ISC marker^8^. Based on Lgr5 reporter allele and CreER lineage tracing assays, Lgr5^+^ cells were first postulated to correspond to crypt base columnar (CBC) cells in the +1 to +3 region^8^. Surprisingly, however, our analyses reveal no statistically significant correlation between *Lgr5* expression and inferred cell potency (Extended Data Fig. 4a-b). Specifically, cells presenting the highest *Lgr5* expression levels were not associated with highest stemness potential; rather, the highest potency was detected in some Lgr5^low^ cells (Fig. 2a, Extended Data Fig. 4c).

**Figure 4.**
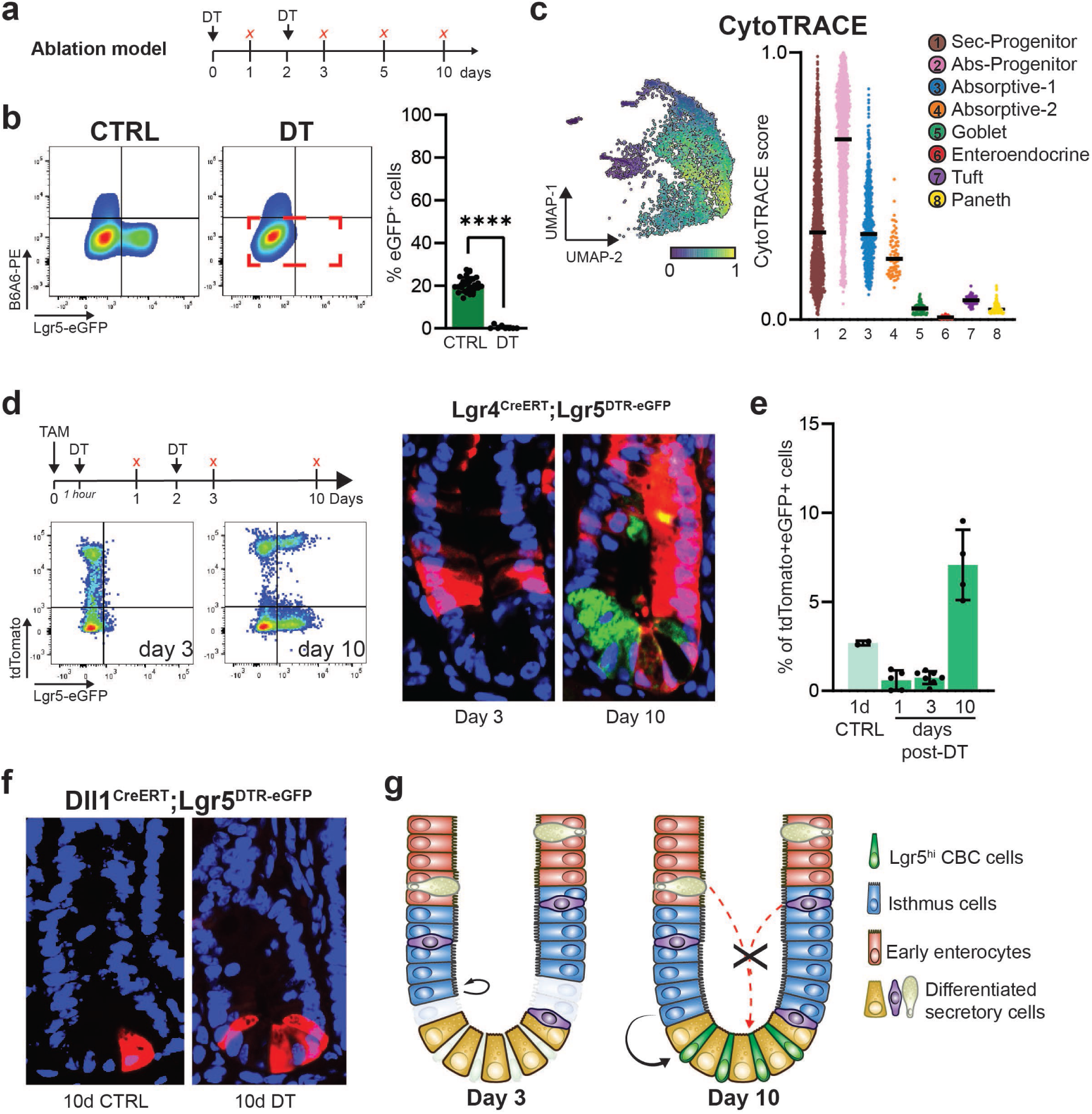
Lgr5neg proliferating cells compensate for loss of *Lgr5* expressing cells: a. Schematic representation of DT based ablation model. **b.** On the left, representative dot-plot of gating strategy adopted for analysis of crypt epithelial cells in CTRL or DT ablated mice (sorted gate in red). On the right, bar-plot showing flow cytometry quantification of Lgr5^DTReGFP+^ cells in control and DT treated crypts (n≥10). **c.** UMAP and violin plots showing CytoTRACE scores in DT treated epithelial cells. **d.** Schematic representation of the model of study together with representative scatter plots for flow-cytometry-based analysis of lineage tracing (Lgr4^CreERT^). On the right, representative images for Lgr4^CreERT^;Lgr5^DTReGFP^ mice lineage tracing upon concurrent DT treatment. **e.** Bar-plot showing quantification of double positive (tdTomato^+^/Lgr5^DTReGFP+^) cells at the indicated time points (Lgr4^CreERT^). (n≥4) **f.** Representative images of Dll1^CreERT^ mice 10 days post TAM induction with or without DT treatment. **g.** Schematic representation of the model proposed, on the left layout of intestinal crypts 3 days post DT treatment, at day 10 (on the right) cells turn back to homeostatic organization. Isthmus cells (blue) expand following loss of Lgr5+ cells, supporting normal tissue turnover. On the other hand, differentiated cells retain low cell potency and do not serve as facultative stem cells. Statistical method: Unpaired t-test, two-tailed; *: p<0.05, **: p<0.01, ***: p<0.01, ****: p<0.01.

To better characterize the *Lgr5* expression pattern, we generated additional scRNA-seq profiles from highly purified crypt epithelial cells harvested from an Lgr5^DTR-eGFP^ mouse^9^ (Fig. 2b). By mapping the *DTR-eGFP* allele on sequencing data, we were able to study expression profiles of both wild type (Wt) and transgenic (Tg) Lgr5 alleles in purified crypt epithelial cells (Fig. 2c). Quantification of the number of cells expressing Wt or Tg alleles revealed virtually complete overlap, confirming that nearly all *Lgr5* expressing cells were also positive for *DTR-eGFP* mRNA(Fig. 2c). Intriguingly, *Lgr5* expression was detected in about 20% of all sequenced crypt cells in our datasets (20.495±1.8, % of total), suggesting a much broader domain of expression then previously appreciated^8^. Flow cytometry analysis of the crypt epithelium consistently revealed an average of 20% Lgr5^eGFP+^ cells (19.96±3.96 % of Epcam^+^ cells), thus confirming the faithful activity of the reporter allele, which was not confined to the CBC cells population (Fig. 2d).

We next analyzed the identity of *Lgr5* expressing cells within our datasets. Similar to other proposed ISC-specific markers^1–4, 6^, Lgr5 expression was broadly detected in multiple intestinal cell types (Extended Data Fig. 4d). Most *Lgr5^hi^* cells were detected in the Sec-Progenitor subpopulation (Extended Data Fig. 4e). In line with this, *Lgr5* expression was found to be associated with the secretory lineage (Fig. 2e, Extended Data Fig. 4f). Moreover, we detected a small fraction of *Lgr5*-expressing cells that were also positive for known secretory markers, such as *Atoh1* and *Dll1*, in agreement with previous studies that have observed direct specification of secretory cells from Lgr5^+^ ones^17, 19^ (Extended Data Fig. 4g-h). Surprisingly, we also observed high levels of *Lgr5* expression in differentiated tuft cells marked by *Dclk1* expression^19, 42^ (Extended Data Fig. 4e,g). Dclk1 staining validated its overlap with Lgr5^eGFP+^ cells, both within the crypt as well as in the villus region (39.2±9.5 % of Dclk1^+^ cells; Fig. 2f, Extended Data Fig. 4i). Moreover, sorted Dclk1^ZsGreen+^ cells^43^ confirmed expression of *Lgr5* and *Ascl2*^44^, both proposed ISC markers, in differentiated tuft cells (Extended Data Fig. 4j-k). Taken together, these data show that *Lgr5* expression is not restricted to intestinal stem and progenitor cells but also spans differentiated secretory subpopulations, including tuft cells.

### Stemness potential exists beyond Lgr5^+^ cells

Lgr5^eGFP-CreERT^ mice show clear labelling of a subset of crypt epithelial cells exhibiting long term lineage tracing, consistent with an ISC^8^. Nevertheless, our analyses indicate that *Lgr5* expression *per se* does not distinguish cells with potency from differentiated ones; rather stemness potential characterize broadly stem and progenitors cells, with highest levels of cell potency detected in some Lgr5^low^ cells (Fig. 2a). In order to investigate the ISC potential of cells other than Lgr5^+^ CBC ones, we studied their capacity to generate organoids *in vitro*^45^. Lgr5^neg^ cells could form organoids (Extended Data Fig. 5a) and enrichment for Ki67^RFP+;^Lgr5^DTReGFP-(neg)^ proliferative cells^46^ increased their efficiency (Fig. 3a); thus, in agreement with our analysis, stemness potential is not restricted to *Lgr5* expressing cells.

**Figure 5.**
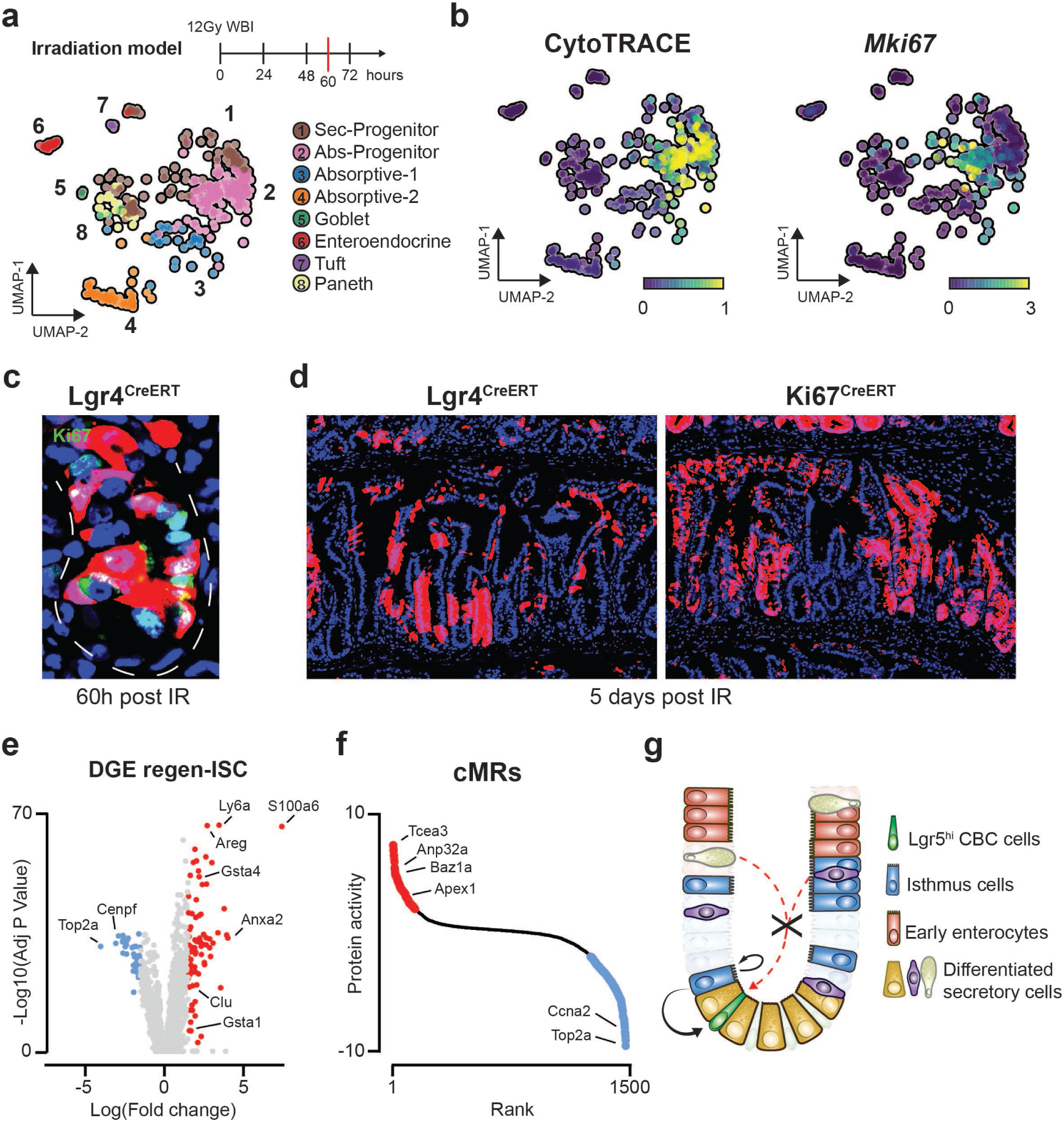
Surviving isthmal stem cells regenerate the intestinal epithelium following IR damage: a. Schematic representation of irradiation damage model, together with UMAP plot showing protein activity-based clustering solution for irradiated crypt epithelial cells analyzed 60 hours post IR exposure. **b.** UMAP plots showing both inferred cell potency through CytoTRACE and *Mki67* expression in irradiated crypt epithelial cells. **c.** Representative image for Ki67 staining 60h post IR, in red tdTomato labelling derived from lineage tracing of Lgr4^CreERT^. **d.** Representative images of Lgr4^CreERT^ and Ki67^CreERT^ linage tracing 5 days post IR. **e.** Volcano plot of differentially expressed genes in regenerating (positive LogFC) vs homeostatic ISC (negative LogFC). **f.** Signature of differentially activated regulatory proteins in ISC resulting from exposure to IR damage. Proteins are ranked (left to right) based on computed protein activity score (NES, normalized enrichment score). **g.** Schematic representation of the model proposed of surviving intestinal cells after IR exposure. Surviving pre-existing stem and progenitors expand and drive tissue repair while differentiated cells are unable to serve as regenerating ISCs.

To clarify if organoid forming capacity reflects functional stemness or a common feature of intestinal epithelial cells, by means of de-differentiation, we employed multiple engineered mouse models that allow discriminating stem and progenitor cells from differentiated cells and evaluated their capacity to generate organoids *in vitro*. To enrich for progenitors^47^ we generated a novel mouse model (Lgr4^CreERT^) that targets inducible Cre to *Lgr4* expressing cells. Consistent with our expectation, following TAM induction, a majority of labelled cells were actively cycling and localized to the crypt isthmus (+4/+12 positions), where proliferation is highest (Fig. 3b, Extended Data Fig. 5b). Similarly, the Ki67^CreERT^ mouse model^48^ confirmed efficient labelling of cycling cells within the crypts and presented a similar spatial distribution (Extended Data Fig. 4b). Over time, both models gave rise to tracing ribbons, which persisted for up to six months, thus demonstrating labelling of some ISCs (Fig. 3c, Extended Data Fig. 5c-d). Conversely, to selectively label differentiated cells we generated a novel mouse model that targeted inducible Cre to *Dll1*-expressing cells (Dll1^CreERT^). Dll1^CreERT^ mice efficiently label committed secretory cells that overlap only minimally with proliferative progenitors and do not give rise to long term lineage tracing (Fig. 3b-c). Moreover, we studied our previously described Dclk1^-ZsGreen-DTR^ mouse line that efficiently labels tuft cells within the intestine (Extended Data Fig. 4j-k)^43^. Notably, as previously reported, differentiated tuft cells do not expand and the majority are lost within one week from labelling (Extended Data Fig. 5e)^49^. In agreement with cell potency inference (Fig. 1d), only stem/progenitor cells could form organoids *in vitro* (Fig. 3d), suggesting that organoids forming capacity reflects stemness competence rather than plasticity, and that exposure to growth factors alone is not sufficient to promote epithelial cells de-differentiation.

These results show that cells other than Lgr5^+^ CBC cells recapitulate functional stemness properties *in vitro.* Intriguingly, recent intravital studies have reported downward migration of cells at the base^50^, suggesting that positional advantage does not constitute a limiting factor for niche competition^51^. Together, these and our observations raise the possibility that cells outside the base may contribute to normal tissue turnover. To determine if isthmus cells can exert downward migration and give rise to CBC cells, we sought to carefully analyze the tracing behavior of Lgr4^CreERT^ and Ki67^CreERT^ mouse lines. Albeit not specific to the crypt isthmus, these mouse models show minimal labelling of the CBC cells’ compartment while including cells capable of long-term lineage tracing (Extended Data Fig. 5b,e), making them an attractive model to test this hypothesis. Quantification of positional labelling over time indicated bi-directional clonal expansion of traced cells from the crypt isthmus, with some invading the CBC cells compartment (Fig. 3e-f, Extended Data Fig. 5f-g). This was further supported by Cre-independent analysis of proliferative cells’ migration (Extended Data Fig. 5h-j). Additionally, analysis of the overlap between labelled cells and Lgr5^DTReGFP^ indicated enrichment for Lgr5^low^ cells in the double positive – tdTomato^+^;Lgr5^DTReGFP+^ - population (Fig. 3g, Extended Data Fig. 5k), supporting the isthmus origin of the expanded clones.

Altogether these analyses indicate that stemness potential is not restricted to *Lgr5*-expressing cells but rather covers broadly the stem and progenitor compartments. Furthermore, they provide evidence that isthmus cells can serve as an ISC compartment and contribute to cellular turnover during intestinal homeostasis.

### Lgr5^neg^ isthmus cells compensate for loss of Lgr5 expressing cells

Our data indicate that alongside Lgr5^+^ cells, other crypt epithelial cells retain functional stem cell properties. This is consistent with multiple studies showing that selective ablation of *Lgr5* expressing cells does not impact normal tissue turnover. In particular, two models of interpretation have been proposed to explain such observations, whereby Lgr5^+^ cells repopulation is driven by either (a) de-differentiation of terminally differentiated cells^10–13^, or (b) activation of a reserve quiescent ISC^7, 9^.

To clarify the dynamics of this process we analyzed the diphtheria toxin (DT) ablation model using the Lgr5^DTReGFP^ mouse line^9^. Flow cytometry analysis confirmed consistent, complete elimination of Lgr5^DTReGFP+^ cells after two doses of DT without compromising the intestinal mucosa; this was followed by the later reappearance of *Lgr5* expressing cells, returning to control values within 10 days (Fig. 4a, Extended Data Fig. 6a). Of note, the disappearance of Lgr5^+^ cells coincided with a marked reduction of Dclk1^+^ tuft cells (Extended Data Fig. 6b), corroborating their expression of *Lgr5* and aligning well with previous reports^52^.

To gain additional insights into the response to Lgr5^+^ cell ablation, we profiled the transcriptome of purified crypt epithelial cells at 24h after two doses of DT; this represents a time point when most Lgr5^+^ cells are ablated (% of Lgr5-eGFP^+^ cells: 0.44±0.85 vs 20.52±1.94 - Fig. 4b) and regeneration has begun. ScRNA-seq profile analysis revealed that, despite marked reduction in *Lgr5* expressing cells, the overall cellular composition remained intact (Extended Data Fig. 6c-d), consistent with the observation that Lgr5^+^ cells are largely dispensable for crypt epithelium regeneration^9^. Importantly, these analyses did not reveal expansion of any novel cell populations that may be ascribed to activation of a reserve ISC. Analysis of inferred cell potency of DT-treated crypt epithelial cells, relative to control, revealed dramatic reduction of cells that comprised the top decile of high potency scored cells (Fig. 4c, Extended Data Fig. 6e), in line with the notion that *Lgr5* is expressed in some stem and progenitor cells. Nevertheless, a consistent fraction of high inferred stemness cells persisted at this time point, further supporting the hypothesis that some Lgr5^neg^ cells possess the potential to serve as ISCs to support tissue regeneration.

Similar to control, following DT treatment high cell potency was confined to stem and progenitor cells, suggesting that ISC potential is also restricted to these populations during regeneration. To test this hypothesis, we first analyzed the behavior of Lgr4^CreERT^ labelled cells following DT treatment. Surviving tdTomato^+^Lgr5^DTReGFP-^ cells could be observed scattered across the crypt epithelium and over time expanded from the isthmus giving rise to new tdTomato^+^Lgr5^DTReGFP+^ cells at day 10 (Fig. 4d-e, Extended Data Fig. 6f-g). Similar findings were obtained when exposing mice to prolonged DT treatment, which revealed consistent lineage tracing of Lgr5^neg^ proliferating cells (Extended Data Fig. 6h). Notably, mice had to be sacrificed by day 9 due to toxicity of DT that was independent of Lgr5^DTReGFP^ expression (Extended Data Fig. 6i).

Next, we tested whether differentiated cells may also participate in Lgr5^+^ cell repopulation alongside proliferating cells, thus reflecting the proposed broad plasticity of intestinal epithelial cells^11^. Upon crosses to Lgr5^DTReGFP^ mice and DT treatment, Dll1^CreERT^ labelled cells did not expand but rather appeared as Paneth cells at day 10, suggesting that committed and/or differentiated secretory cells do not contribute to Lgr5^+^ cell repopulation (Fig. 4f, Extended Data Fig. 7a-b). A possible explanation for the discrepancy with previous reports^10, 12^ is the higher specificity of our newly generated mouse line for committed secretory cells relative to the previously reported Dll1^eGFPCreERT^, which showed lineage tracing events also in homeostatic conditions^12^. Indeed, low *Dll1* expression can be detected broadly within the stem/progenitor compartment, and this appears true also for other proposed markers of differentiation including *Alpi*, *Fabp2*, and *Krt20* (Extended Data Fig. 7c-d).

Taken together, these data show that, upon loss of *Lgr5* expressing cells, Lgr5^neg^ isthmus cells have the capacity to act as ISCs and to support tissue turnover. Furthermore, they indicate that regenerative potential is restricted to stem and progenitor cells, and that neither de-differentiation nor activation of a reserve quiescent-ISC are the source of Lgr5^+^ cell repopulation in this model (Fig. 4g).

### Surviving isthmus cells regenerate the intestinal epithelium following IR damage

As discussed, analysis of CTRL and DT-ablated crypt epithelial cells revealed neither the presence of a putative reserve ISC population nor any sign of active de-differentiation. Rather, it supports a model where some Lgr5^neg^ cells retain ISC potential and sustain intestinal tissue turnover. To determine whether this may be applicable to other intestinal injury models, we assessed whether intestinal regeneration after lethal irradiation (IR) follows similar cellular dynamics. Damage due to high dose IR exposure can induce the gastrointestinal syndrome, with highly proliferative cells and *Lgr5*^+^ CBCs representing subpopulations more likely to be lost due to irreversible IR damage^8^.

To pinpoint the identity of cells with regenerative potential, we studied crypt epithelial cell composition after exposure to 12 Gy whole body IR (WBI), by scRNA-seq analysis. First, to identify the earliest stage of intestinal regeneration, we characterized the dynamics of intestinal proliferation following IR damage (Extended Data Fig. 8a). We postulated that at this time point, regenerating stem cells would be undertaking their first or second round of cell divisions, marked by an increase in Ki67 labeling. We identified 60 hours post IR as the earliest time point when proliferation increases, reflecting the first regenerative wave and the focal point for our analysis (Fig. 5a).

Following IR damage, surviving cells aligned well with previously identified clusters and no novel subpopulations emerged (Fig. 5a). As expected^8^, proliferative progenitors as well as *Lgr5* expressing cells were largely depleted at this time point (Extended Data Fig. 8b). Similar to the ablation model, surviving differentiated cells retained low levels of inferred cell potency (Extended Data Fig. 8c), and evaluation of Dll1^CreERT^ or Dclk1^CreERT^-driven lineage tracing corroborated the absence of any participation by differentiated secretory cells in intestinal regeneration (Extended Data Fig. 8d).

In line with our hypothesis, we observed a small fraction of surviving cells that were characterized by high levels of inferred cell potency and expression of proliferative markers (Fig. 5b), thus identifying them as the most suitable regenerating ISCs candidates. Regenerating and homeostatic cells with high cell potency showed a high degree of overlap (Extended Data Fig. 8b), suggesting that regenerating ISCs may correspond to surviving isthmus cells. To clarify the nature of the regenerating ISCs, we lineage traced intestinal proliferating cells following high doses of IR. Sixty hours after IR exposure, surviving tdTomato^+^ cells (Lgr4^CreERT^) could be observed within the damaged epithelium (Fig. 5c, Extended Data Fig. 8e), thereby demonstrating that not all proliferative cells are lost to IR induced damage. Moreover, Ki67 staining and lineage tracing analyses at 5 days post IR confirmed active participation of surviving isthmus proliferative tdTomato^+^ cells in intestinal regeneration (Fig. 5c-d).

We next sought to identify the regulatory programs that characterize regenerating ISCs. For this purpose, we searched for features unique to cells with highest inferred cell potency pre- and post-IR (Fig. 5e-f, Extended Data Fig. 8f, Extended Data Table 9-10). Pathway analysis showed that regenerating ISCs activate cell cycle damage checkpoints and upregulate multiple damage response factors^53–55^, reflecting the profound damage generated by IR exposure. This was further corroborated by analysis of the regulatory proteins that define the regenerating ISC compartment (Fig. 5f). Indeed, Apex1 and Baz1a, both known to be involved in DNA damage response^56, 57^, were among the most statistically significant activated proteins. Moreover, regenerating ISCs expressed high levels of *Ly6a* (Sca1), as well as *Areg* and other proposed intestinal regeneration markers^58^ (Fig. 5e). Interestingly, we observed upregulation of *Clu* (Fig. 5e, Extended Data Fig. 8g), whose expression has been proposed as specific to a quiescent radio-resistant stem cell population (“revival ISC”^7^).

These results raised the possibility that regenerating ISCs may represent an expanding reserve stem cell subpopulation. To clarify the nature of these cells, we analyzed *Clu* expression in traced intestinal proliferating cells (Ki67^CreERT^) pre- and post-IR damage. Results showed *Clu* upregulation in surviving tdTomato^+^ cells sixty hours post IR exposure (Extended Data Fig. 8h), providing evidence that following IR damage, the proposed ‘revival’ ISCs correspond to surviving cycling cells rather than a separate quiescent stem cell population. Lastly, although we were unable to discriminate the relative contribution of pre-existing stem cells or early progenitors, our analyses strongly suggest that the potency to serve as regenerating ISCs is restricted to these populations (Fig. 5G).

## Discussion

Here we provide an unbiased characterization of intestinal crypt epithelial cells in homeostasis and regeneration. We find that stemness potential exists beyond the CBC cells compartment in the crypt isthmus, the region previously proposed to accommodate transit-amplifying progenitors. Furthermore, we provide evidence that intestinal regeneration is driven by surviving stem and progenitor cells.

Our optimized isolation protocol—using the B6A6 Ab^24^ to exclude villi cells—ensured high enrichment of the crypt epithelium and reproducibility across all high-throughput datasets, especially for the injured intestine where loss of tissue integrity greatly affects the isolation strategy. Furthermore, our computational pipeline integrated multiple algorithms to unbiasedly recover cell identities and elucidate their hierarchical organization based on single cell transcriptomic and epigenetic profiles. Compared to previous studies, the results obtained show a high degree of similarity^16, 17^ yet provide improved molecular resolution of intestinal stem and progenitor cells. Moreover, the proposed regulatory network-based classification schema offers novel insights on mechanistic determinants of intestinal epithelial cell subpopulations. Intriguingly, many of the identified stem cell regulatory factors align well with previous reports^19^, confirming their findings and suggesting that such signatures may be useful to study the role of novel factors in defining specific intestinal lineages.

Surprisingly, *Lgr5* expression did not correlate with inferred cell potency. *Lgr5*^hi^ cells were largely found in the clusters associated with secretory-biased progenitor cells, as previously suggested for some intestinal Lgr5^+^ cells^17, 59^, and in line with findings in the stomach, where high levels of *Lgr5* can be detected in differentiated chief^60^ and antral basal secretory cells^61^.

Moreover, Lgr5 expression *per se* does not discriminate between cells with potency from differentiated ones. Stemness potential is found well outside the CBC cell compartment, and isthmus cells can participate in normal tissue homeostasis. Importantly, these observations resonate well with our and others previous reports^1, 4–6^ and highlight similarities with the gastric mucosa where contribution of cells from the isthmus region in tissue cellular turnover has been described^62^. Thus, suggesting that proposed competition dynamics^51^ may involve broadly CBC and isthmus cells compartments.

Cells with the highest inferred cell potency were well segregated from Lgr5^hi^ cells and detected at the boundary between the progenitors clusters (Sec-Progenitor and Abs-Progenitor). This suggest a distinction between highest inferred stemness potential and proposed relative contributions to long term lineage tracing. While some of the applied computational methodologies may suffer of intrinsic biases^20^, this may indicate that selective lineage engagement biases intestinal epithelial cells for long term retention.

Furthermore, the difficulty in segregating cells with highest potency from early progenitors suggests their high-degree of similarity and may indicate their co-existence in closely comparable cellular states. Analysis of the factors that best associate with inferred cell potency revealed multiple chromatin remodelers, including Atad2, which has been recently shown to determine cell potency in the skin^34^, as well as other factors previously suggested to regulate intestinal stemness^36, 37^. Whereas, similarly to Lgr5, other proposed markers for ISC, such as Ascl2^63^, also failed to correlate with inferred cell potency and were rather broadly expressed within the secretory lineage.

In addition to characterizing homeostatic crypt cell composition, we also analyzed intestinal regeneration following Lgr5^+^ cell ablation and IR damage. In line with previous observations^7, 9^, both models mostly eliminated *Lgr5* expressing cells without evoking major composition changes within the intestine, especially for the DT ablation model. Notably, our results indicate that cells with high inferred cell potency persist within the intestine and align well with the proposed homeostatic isthmus compartment. Furthermore, our findings highlight that differentiated cell types retain low cell potency states in both injury models and do not appear to acquire features of stemness. In fact, differentiated secretory cells marked by high levels of *Dll1* or *Dclk1* expression do not participate in intestinal regeneration, whereas surviving isthmus cells do. While the mouse models studied do not allow for the discrimination of the relative contribution between surviving ISCs or early progenitors, they provide evidence that the potential to regenerate is restricted to these populations. In addition, when we analyzed the changes in the transcriptional profile of ISCs following IR damage, we detected a specific signature of regenerating cells^58^. This included upregulation of *Ly6a* (Sca-1) and *Areg* for example, and was accompanied by increased activation of several damage response factors, such as Baz1a and Apex1^56–58, 64^. Intriguingly, we noted upregulation of *Clu*, an additional target of YAP signaling^65^, indicating that previously proposed ‘revival’ stem cells^7^ correspond to surviving ISCs and early progenitors rather than a distinct quiescent stem cell population.

In summary, our unbiased analyses suggest a new regulatory model whereby isthmus cells possess functional stem cells properties and participate both in homeostasis and regeneration. Previous findings suggesting the presence of a reserve stem cell or extensive cellular plasticity likely reflected overlap of inducible Cre driver with the isthmus compartment. Our finding of broad expression by intestinal progenitors of multiple genes used for lineage tracing can help reconcile previous reports^12, 13^. Future studies will need to focus on the regulation of these early events and the key niche signals which maintain stem cell renewal within the intestinal isthmus.

## Author contribution

EM, AV, AC, TCW conceived, designed the study and wrote the manuscript. EM performed most of the experiments. AV performed the computational analyses. YO performed *in vitro* experiments. WK generated and characterized Dll1 transgenic mice. MM, HN, BB, JL, LBZ participated in performing the experiments. MHW provided the B6A6 antibody. LL provided critical conceptual input and manuscript revisions. KSY, CWC, and CG participated in the discussion. TCW secured the funding.

## Acknowledgments

This research was funded in part through the NIH/NCI Cancer Center Support Grant P30CA013696 and used the resources of the Herbert Irving Comprehensive Cancer Center Flow Cytometry Shard Resources, Molecular Pathology/MPSR, Genomics and High Throughput Screening, as well as the Genetically Modified Mouse Model Shared Resource (GMMMSR). This research was also supported by the Columbia University Digestive and Liver Disease Research Center (CU-DLDRC) grant 1P30DK132710. This work was supported by grants from the NIH/NCI including UO1DK103155, R35CA210088, RO1NK128195 to TCW; as well as a NCI Outstanding Investigator Award (R35 CA197745) and two NIH Shared Instrumentation Grants (S10 OD012351 and S1 0OD021764), all to AC. AV is supported by a U.S. Department of Defense Early Investigator Research Award (W81XWH19-1-0337) and an Early Career Development Pilot Award NIH/NCI Cancer Center, funded through the Cancer Center Support grant, P30CA013696. We are grateful for the support we received by Columbia University shared resources and we would like to thank Sun Dajiang “Kevin” (Molecular Pathology/MPSR), Kissner, Michael (Columbia Stem Cell Initiative Flow Cytometry core facility), Erin Bush (Genomics and High Throughput Screening), and Lin Chyuan-Sheng “Victor” (Genetically Modified Mouse Model), for their incredible expertise and help in this project. We thank Nicoletta Barolini (Columbia University) for designing the graphical models.

## Extended Data Figures and Methods

**Extended Data Fig. 1:**
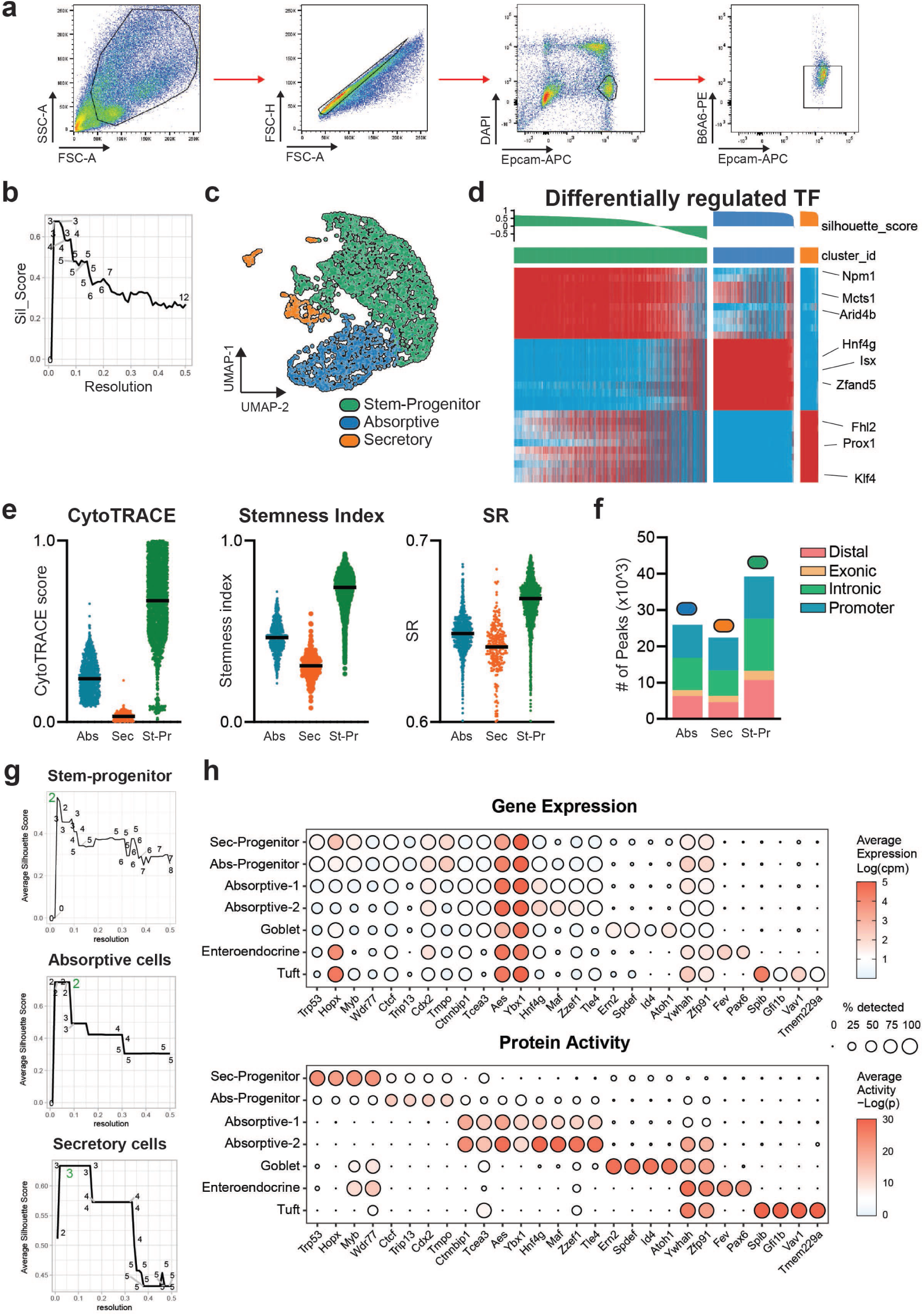
**a.** Gating strategy used for flow cytometry-based sorting of crypt epithelial cells. **b.** Showing Louvain’s clustering resolution parameter optimization between 0.01 and 0.5. On the y-axis showing the average silhouette score for each clustering solution of crypt epithelial cell regulatory protein activity as identified using the relative resolution parameter shown on the x-axis. This optimization strategy yielded to n=3 clusters. **c.** UMAP plot showing protein activity-based clustering solution. **d.** Heatmap showing top differentially activated regulatory proteins (activated proteins in red color). Silhouette scores for individual cells shown on the top of the heatmap. Cells in each cluster are sorted by silhouette score. **e.** Violin plots showing CytoTRACE, Stemness index, and Signaling Entropy (SR) scores computed for individual cells. **f.** Bar plot showing number of identified accessible peaks per cluster based on scATACseq. **g.** Louvain’s clustering resolution parameter optimization as 1-step iterative sub-clusters identification applied to each one of the n=3 clusters identified on crypt epithelial cells (i.e., stem-progenitors, absorptive and secretory cells). In green highlighted the optimal number of clusters for each iteration. **h.** Showing dot-plots of top 4 markers for each cluster (rows) as identified from protein activity-based clustering. Panel on the top shows gene expression as log(cpm), while panel on the bottom shows for the same markers the corresponding protein activity as -log10(p).

**Extended Data Fig. 2:**
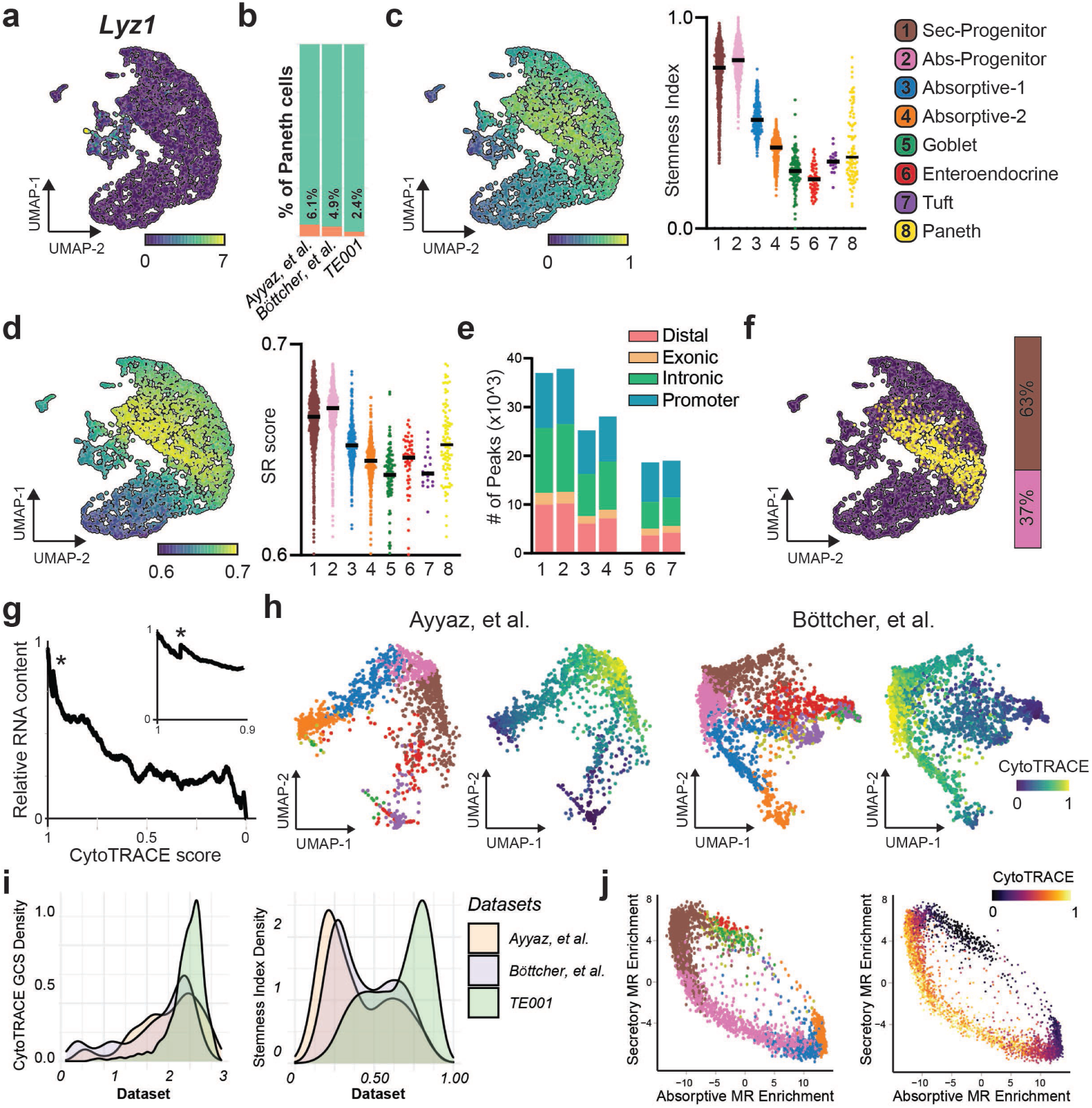
**a.** UMAP plot showing expression domain for *Lyz1* on crypt epithelial cells. **b.** Bar-plot showing quantification of Paneth cells (in orange) across the datasets tested. Candidate Paneth cells were identified based on the concurrent expression of *Lyz1*, *Mmp7*, *Dll4*, and *Nupr1*. **c-d.** UMAP and Violin plots for stemness-index (c.) and entropy (d.) scores computed for individual crypt epithelial cells. **e.** Bar plot showing number of accessible peaks per cluster together; note cluster 5 consist of very few cells and was therefore excluded from further analyses. **f.** UMAP plot showing top 10% cells based on CytoTRACE score, on the right bar-plot showing cluster contribution to the top 10% population. **g.** Plot showing individual cells’ values of Relative RNA content and Cytotrace scores. In small, showing only the top 10% of cells based on CytoTRACE levels. Black star indicates the ‘valley’ as previously described^1^. **h.** UMAP plots of homeostatic crypt epithelial cells from Ayyaz et al. and Bottcher et al. analyzed at protein activity, together with CytoTRACE analysis in the two datasets. Clusters are assigned to each cell by performing enrichment analysis on MRs signatures as identified by us. **i.** Density plots showing distribution of CytoTRACE GCS^1^ and Stemness-index scores^2^ for the three datasets tested. **j.** Showing homeostatic crypt epithelial cells analyzed at protein activity on a scatter plot where each axis represents the extent of enrichment for MRs candidate drivers of absorptive or secretory lineages respectively; on the right, computed CytoTRACE scores are overlaid.

**Extended Data Fig. 3:**
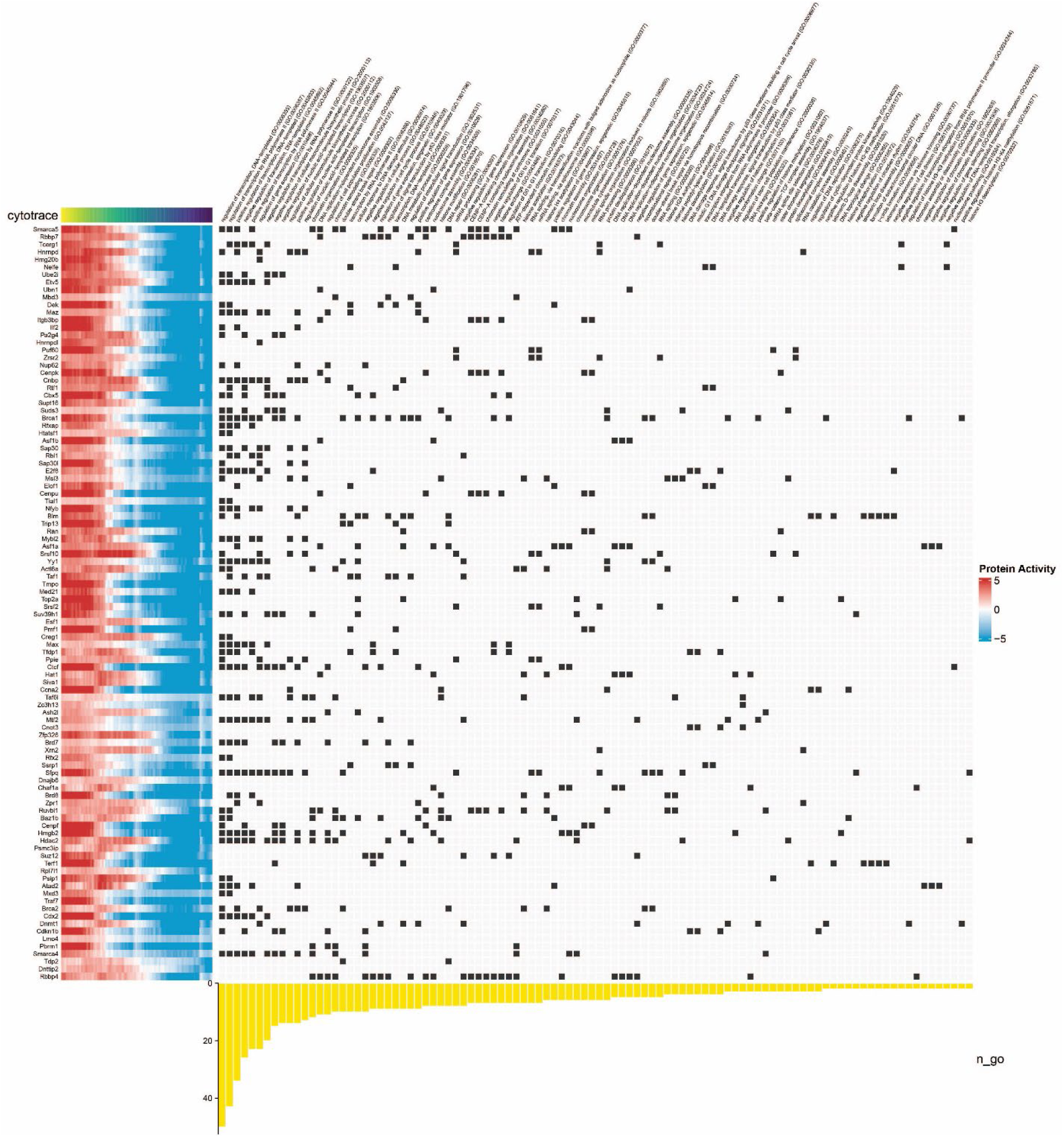
On the left, strip plots showing inferred activity of the 100 CytoTRACE score-based top-correlating transcriptional regulators. On the right, top GO-associated terms for each regulatory protein, using ‘Molecular Function” as query.

**Extended Data Fig. 4:**
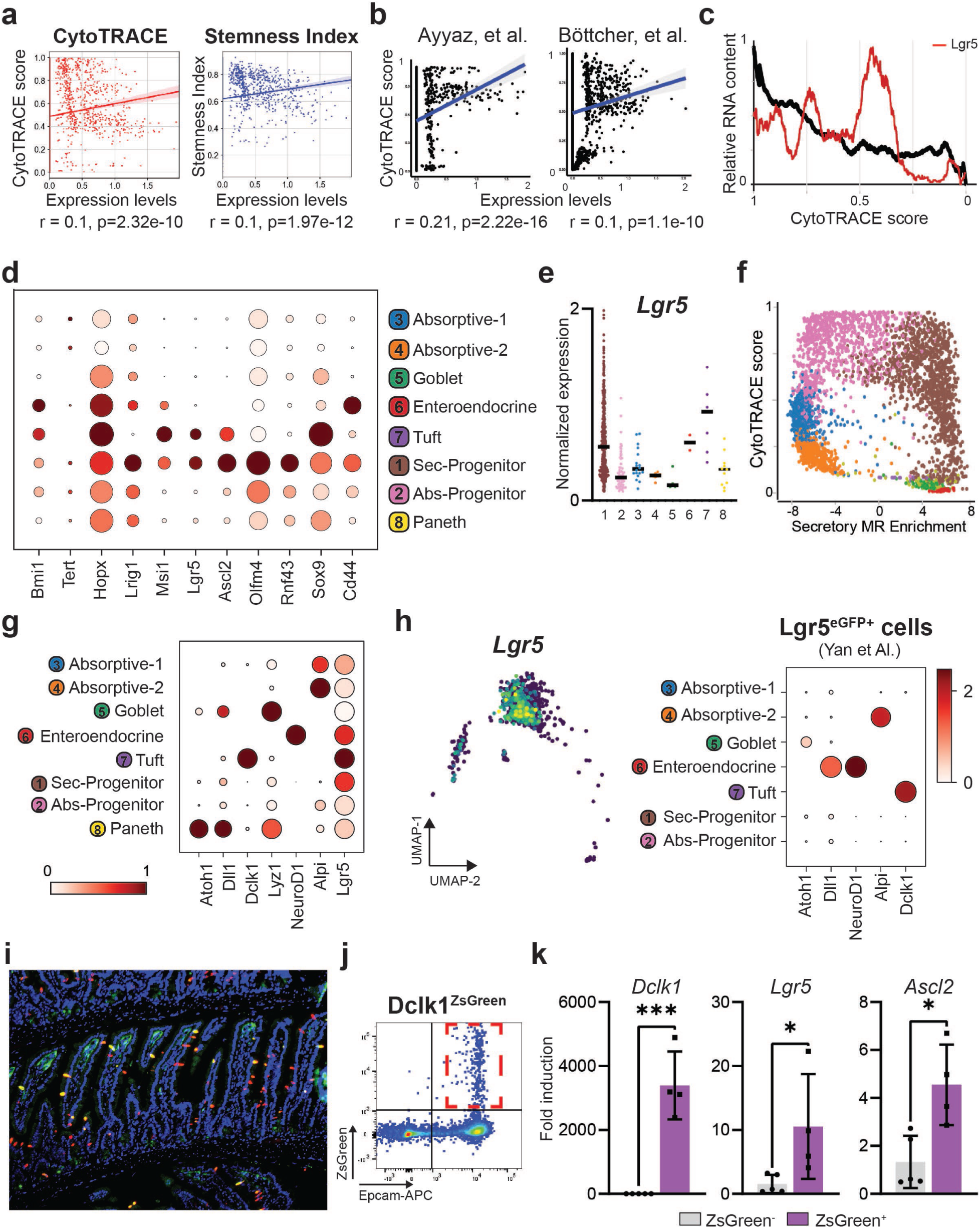
**a.** Scatter plots showing Pearson’s correlation analysis between *Lgr5* expression and inferred cell potency, as assessed by CytoTRACE (left) and Stemness Index (right). **b.** Scatter plots showing Pearson’s correlation analysis between *Lgr5* and CytoTRACE scores in the analyzed datasets. **c.** Plot showing individual cells’ values of Relative RNA content and Cytotrace scores with Lgr5 expression distribution overlaid in red. **d.** Dot-plot showing expression of selected genes in crypt epithelial cells. **e.** Violin-plot showing expression levels of *Lgr5* in the identified clusters (only Lgr5^+^ cells are shown). **f.** Scatter-plot where inferred cell potency (CytoTRACE, y-axis) is presented together with enrichment scores for secretory (x-axis) lineage. Cells are colored according to clustering solution. **g.** Dot-plot showing expression of known markers of differentiation in Lgr5 expressing cells. (note that only Lgr5^+^ cells are shown) **h.** Analysis of Lgr5^eGFP+^ sorted cells - Source data: Yan KS et al.^3^. Left: UMAP distribution plot showing expression of *Lgr5*; right: dot-plot showing expression of markers of differentiated cells. **i.** Representative image of low power magnification view of Dclk1 (Red) and Lgr5-eGFP (green) in intestinal jejunum of a Lgr5^DTR-eGFP^ mouse. **j.** Sorting strategy for the isolation of Dclk1^ZsGreen+^ cells (Red gate). **k.** Bar-plots showing expression levels of *Dclk1*, *Lgr5*, and *Ascl2* in sorted Dclk1^ZsGreen+^ cells. (Expression presented as fold induction relative to the negative population) (n=4). Statistical method: Unpaired t-test, two-tailed; *: p<0.05, **: p<0.01, ***: p<0.01, ****: p<0.01.

**Extended Data Fig. 5:**
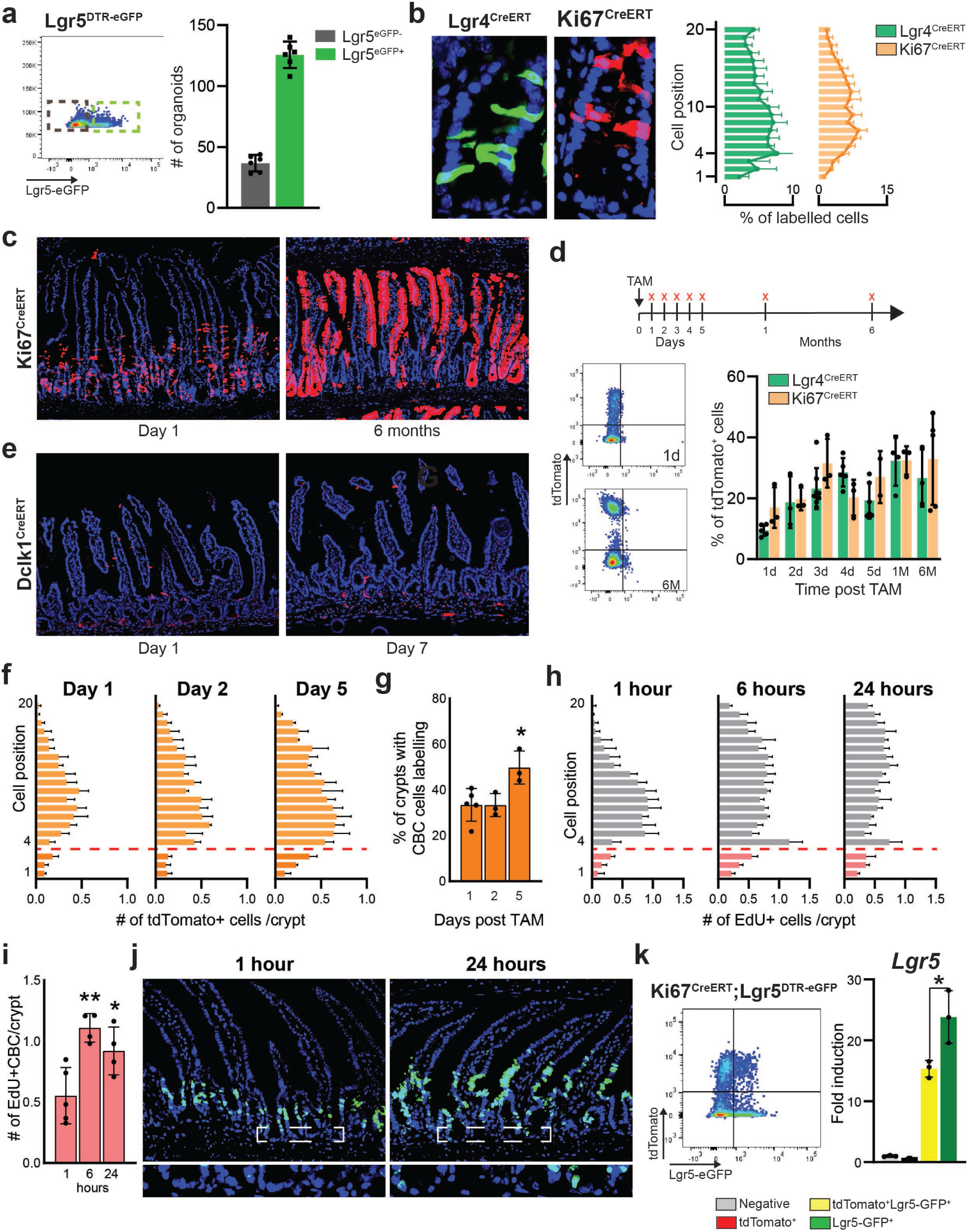
**a.** Representative scatter-plot for flow cytometry based sorting of Lgr5^DTReGFP^ mice. On the right, bar-plot showing quantification of the number of organoids generated in the isolated cells fractions. **b.** Representative images together with bar-plots showing position based counting of labelled cells 24h post TAM administration in Lgr4^CreERT^ and Ki67^CreERT^ mice. (n=3) **c.** Representative images for Ki67CreERT lineage tracing 1 day and 6 months post TAM induction. **d.** On top, schematic representation of the model of study, red crosses indicate time-points of analysis. Bottom, representative scatter plots for flow-cytometry-based analysis of lineage tracing (Lgr4CreERT) together with bar-plot showing flow cytometry-based quantification of tdTomato+ cells at indicated time points (n≥3). **e.** Lineage tracing analysis of Dclk1^CreERT^ mouse model. At day 1 scattered tdTomato^+^ tuft cells can be observed, at day 7 post TAM most tdTomato^+^ cells are washed out. **f.** Bar-plots showing quantification of traced cells in Ki67^CreERT^ mice based on cell position within the crypt. Red dotted line used to mark the boundary between CBC and isthmus cells (n=3). **g.** Bar-plots showing percentage of crypt-units with at least one labelled CBC cell in Ki67^CreERT^ mice at indicated time points. (n=3) **h.** Bar-plots showing quantification of EdU-labelled cells based on cell position at the indicated time points. Red dotted line used to mark the boundary between CBC and isthmus cells (n=3). **i.** Bar-plot showing average number of EdU labelled-CBC cells at indicated time points. **j.** Representative images of EdU staining in the mouse intestine. Below, enlargement of the highlighted region to show absence/presence of CBC labelling. **k.** Flow cytometry analysis of overlap between Ki67 labelled cells (24h post TAM) and Lgr5^eGFP+^ cells; together with bar plot showing expression levels of *Lgr5* in single sorted cells, groups labelled in the panel. (n=3). Statistical method: Unpaired t-test, two-tailed; *: p<0.05, **: p<0.01, ***: p<0.01, ****: p<0.01.

**Extended Data Fig. 6:**
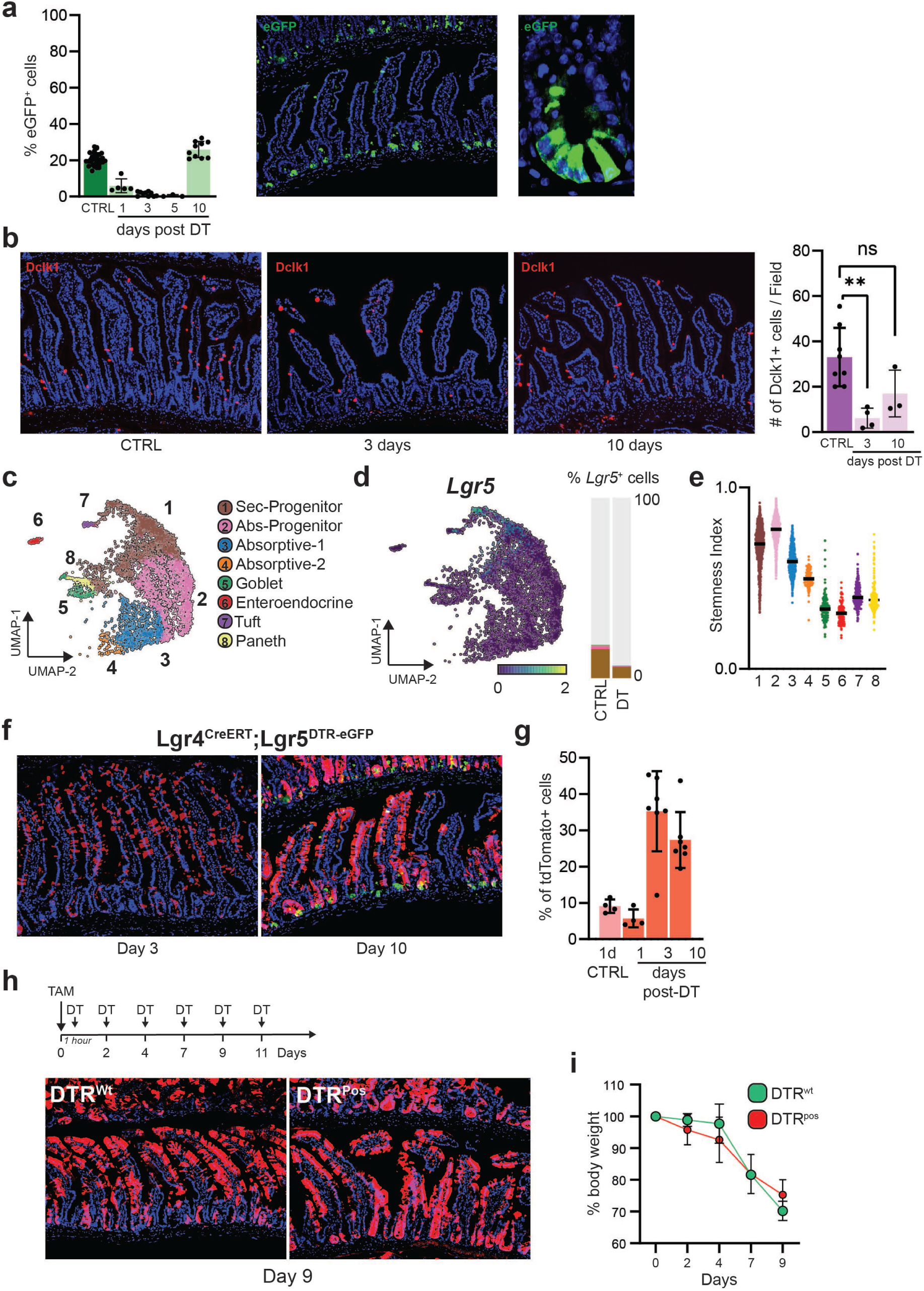
**a.** Flow cytometry quantification of Lgr5^eGFP+^ cells in purified crypts (n≥3). On the right, representative images of Lgr5^eGFP^ 10 days after the first dose of DT. **b.** Representative images for Dclk1 staining of intestine following DT treatment together with bar plot showing quantification of the number of Dclk1+ cells at indicated time points (n≥4). **c.** UMAP plot showing recovered clusters in DT treated epithelial cells. **d.** UMAP plot showing expression levels of *Lgr5* in the DT treated dataset, on the right bar-plot showing % of detected expressing cells relative to control. Cells are colored based on clusters. **e.** Violin-plot showing computed stemness-index for individual DT-treated crypt epithelial cells. **f.** Representative images for Lgr4^CreERT^;Lgr5^DTReGFP^ lineage tracing at day 3 and 10 post DT exposure. **g.** Bar-plot showing percentage of total tdTomato^+^ cells at indicated time-points (Lgr4^CreERT^, n≥4). **h.** Schematic representation of the model used. TAM was administered 1 hour prior the first DT injection. After this DT was injected every two days. Below, representative images of intestinal sections for Ki67^CreERT^ lineage tracing at day 9. **i.** Plot showing percentage of body weight loss recorded over the course of the experiment. No significant differences could be detected between the groups (n=3). Statistical method: Unpaired t-test, two-tailed; *: p<0.05, **: p<0.01, ***: p<0.01, ****: p<0.01.

**Extended Data Fig.7:**
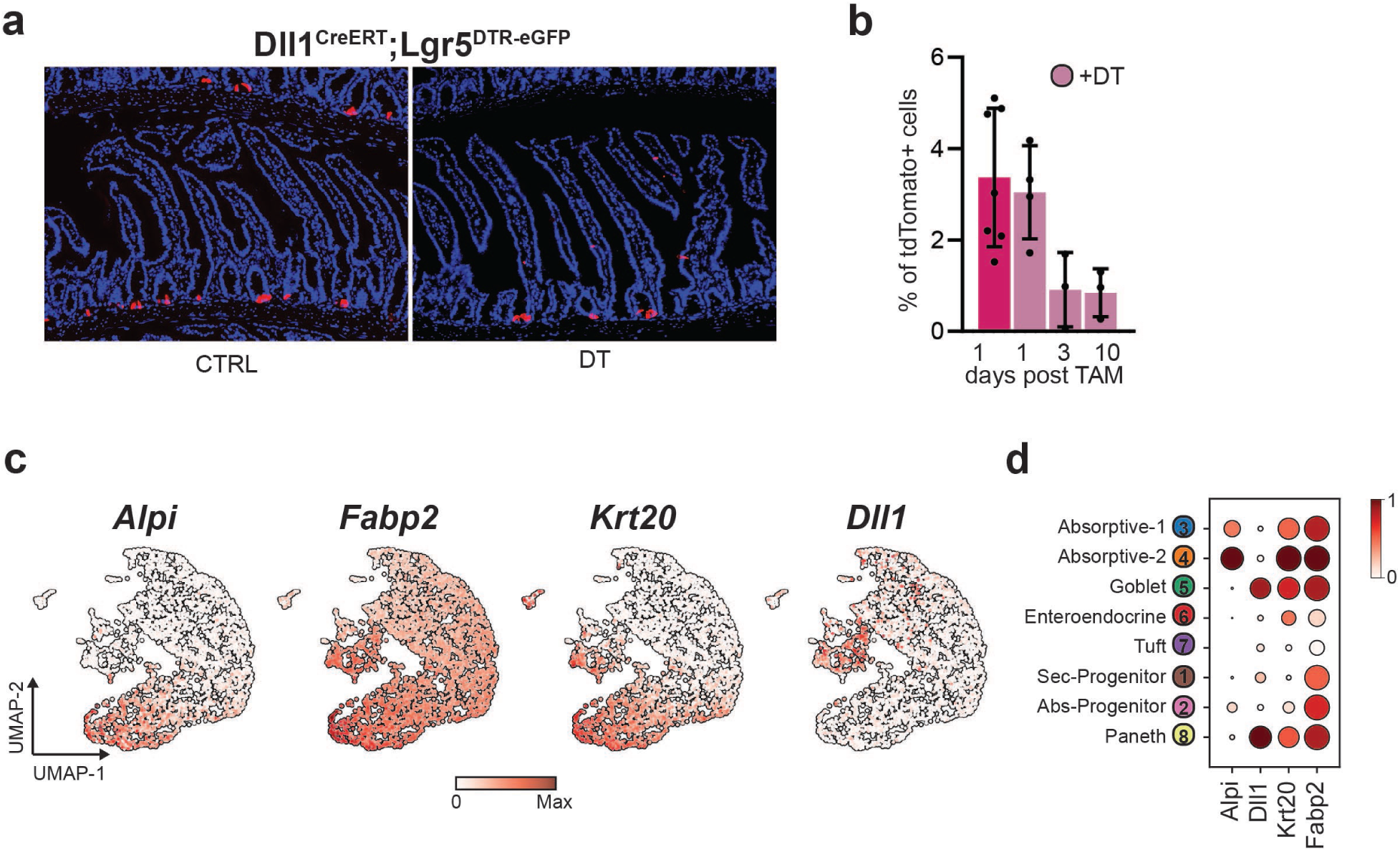
**a.** Representative images of Dll1CreERT mice 10 days post TAM induction with or without DT treatment. **b.** Bar plot showing flow-cytometry based quantification of (Dll1^CreERT^) tdTomato^+^ cells at indicated time points. (n≥3). **c-d.** UMAP and dot-plots showing expression levels of the indicated genes in crypt epithelial cells.

**Extended Data Fig. 8:**
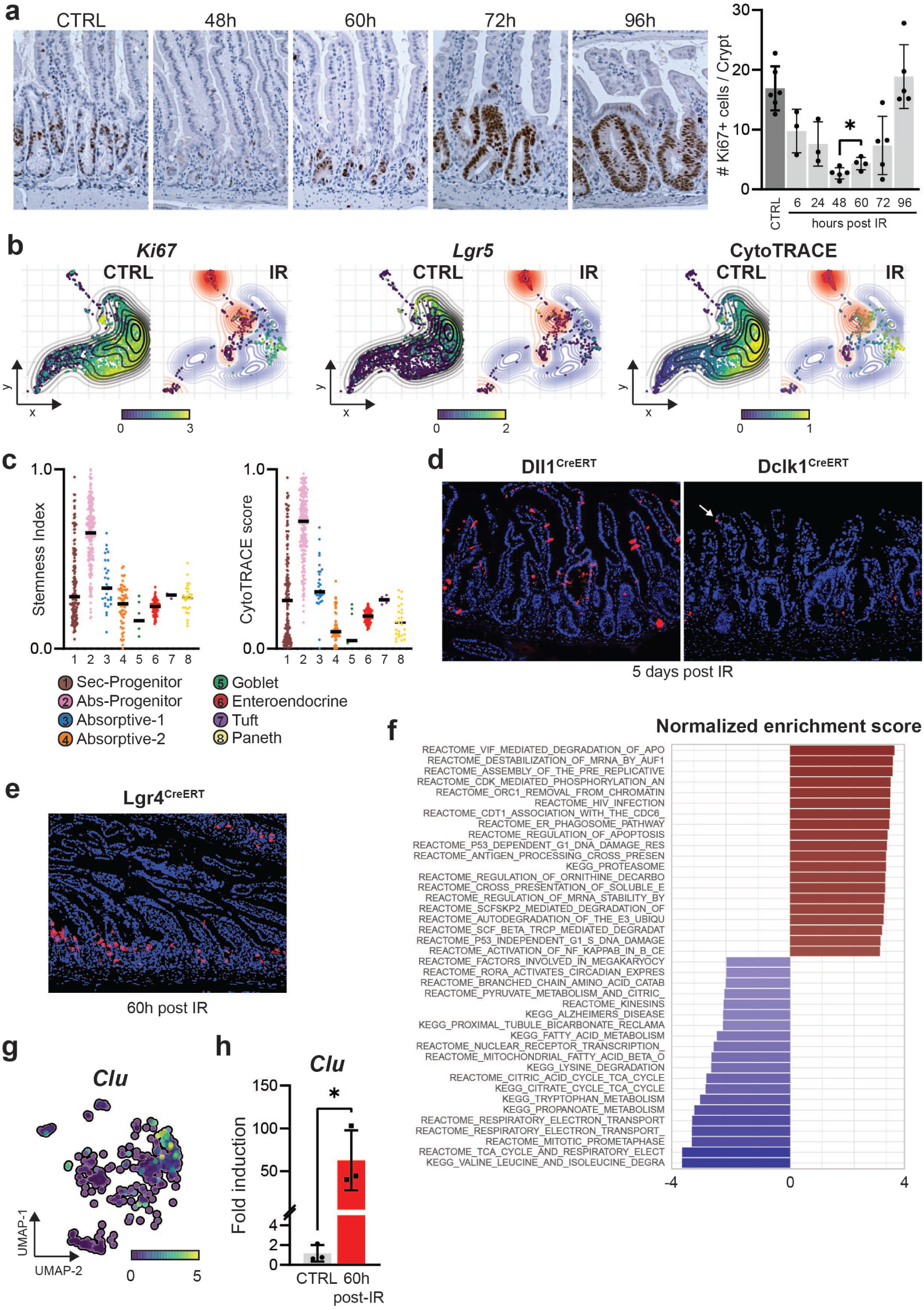
**a.** Immunohistochemistry analysis for Ki67 following IR damage; on the right, bar plot showing quantification of the number of Ki67^+^ cells at indicated time points. (n≥3) **b.** Differential Kernel density estimation (KDE) analysis of irradiated crypt epithelial cells, using a bootstrapped null model for the control. Showing expression for Ki67 (Left) and Lgr5 (middle), as well as computed CytoTRACE scores (right). **c.** Bar-plots showing computed CytoTRACE as well as Stemness Index scores in irradiated crypt epithelial cells. CytoTRACE scores are presented relative to control dataset. **d.** Representative images showing tracing of tdTomato^+^ cells (Dll1^CreERT^ and Dclk1^CreERT^ mouse lines) five days post IR damage. **e.** Representative image of Lgr4^CreERT^ mice 60 hours after IR damage. **f.** Bar-plot showing gene expression-based pathway enrichment analysis**. g.** UMAP plot showing *Clu* expression in irradiated crypt epithelial cells. **h.** qPCR analysis of *Clu* expression in sorted tdTomato^+^ (Ki67^CreERT^ mice) of CTRL and 60h following IR. (n=3). Statistical method: Unpaired t-test, two-tailed; *: p<0.05.

## Extended data tables

**Table 1:** Mouse models used in this study

**Table 2:** Reagents and antibodies

**Table 3:** Gene expression 3-cluster Solution

**Table 4:** Protein activity 3-cluster Solution

**Table 5:** Gene expression 8-cluster Solution

**Table 6:** Protein activity 8-cluster Solution

**Table 7:** Protein activity vs CytoTRACE score correlation

**Table 8:** Gene expression vs CytoTRACE score correlation

**Table 9:** Gene expression signature (IR vs CTRL)

**Table 10:** Protein activity signature (IR vs CTRL)

## Methods

### Animal studies

All animal studies were carried out in compliance with the National Institutes of Health guidelines for animal research and approved by the Institutional Animal Care and Use Committee of Columbia University.

All transgenic mouse lines used in this project are listed in (Extended Data Table 1) together with their respective Tamoxifen (TAM) (Sigma, T5648) dosage. Lgr4-DSRED2-CreERT Knock-in and Dll1-ZsGreen-CreERT BAC mouse models were generated here at Irving Cancer Research Center (The Genetically Modified Mouse Models Core) using standard recombineering strategies. Inducible Cre lines were crossed with R26-tdTomato (Strain #:007909), R26-ZsGreen (Strain #:007906), or R26-Confetti (Strain #:017492) reporter allele purchased from Jax (The Jackson Laboratory). TAM was dissolved in corn oil and administered 200uL per oral gavage at indicated time points. For multicolor tracing analysis, using the R26-Confetti allele, a single dose of 6mg/mouse (TAM) was administered. BrdU (Biolegend, 423401) was injected i.p. at 50mg/Kg 6 hours before harvesting. EdU (Sigma, 900584) was injected i.p. at 50mg/Kg, chase time indicated in the figure panels. Diphtheria toxin (DT -EMD Millipore, 322326) was injected i.p. at 50ug/Kg as previously described^4^. For irradiation studies mice received a single dose of 12Gy whole body irradiation using Mark I Cesium-137 based gamma-ray irradiator (J.L. Shepherd & Associates, San Fernando, USA).

### Crypt epithelial cell preparation and staining

All reagents and antibodies are listed in Extended Data Table 2. 10 cm of intestinal Jejunum was harvested and cut in small pieces, EDTA (Invitrogen, 15575020) based dissociation was performed as previously described ^5^ with minor modifications. Following EDTA dissociation crypts were incubated at 37°C for 7 minutes in TrypLE Express (Thermo Fisher Scientific, 12-604-021) and DNase I (Thermo Fisher Scientific, 10104159001) in order to obtain a single cell suspension. Following enzymatic digestion cells were resuspended in Advanced DMEM/F12 (Gibco, 12634-010) supplemented with GlutMAX (Thermo Fisher Scientific, 35050061), HEPES (Thermo Fisher Scientific, 15-630-080), Antibiotic-Antimycotic (A/A - Fisher Scientific, 15240062), and 10% FBS (Gemini Bio-Products, 900-108). Cells were stained for 20 minutes Epcam-APC (1:200, BioLegend 118214), washed and re-suspended in media containing DAPI (1:10’000, BD Biosciences 564907). For B6A6^6^ staining, cells were incubated with primary antibody for 30 minutes, following conjugated secondary Ab for 30 minutes (1:200, BioLegend 405406), washed and re-suspended in media containing DAPI.

### Flow cytometry analysis and cell sorting

All flow cytometry analyses were performed using a LSRII Fortessa (CCTI Flow core) and results were analyzed using FlowJo. Cell sorting experiments were performed in the Columbia Stem Cell Initiative Flow Cytometry core facility at Columbia University Irving Medical Center under the leadership of Michael Kissner, using a BD FACS Aria II. All flow cytometry quantifications are presented as percentage of Alive/Epcam^+^ cells following standard gating strategy (Cells/SingleCells/DAPI^-^Epcam^+^).

### *In vitro* organoids

Single sorted epithelial cells were resuspended in MEDIA and counted under the microscope. Appropriate volume of cell suspension was mixed to GFR Matrigel (Corning 356231) and plated in a 24-wells plate as 25uL domes, 5000 cells/dome were seeded in all the experiments. ENR media (Advanced DMEM/F12, GlutMAX, HEPES, A/A, B27 (Thermo Fisher Scientific, 17504044), N-2 supplement (Fisher Scientific, 17502048), N-Acetyl-L-cysteine (Sigma-Aldrich, A9165), EGF (Thermo Fisher Scientific, PMG8043), Noggin (PeproTech, 250-38), Rspo1 (R&D Systems, 3474-RS-050) supplemented with CHIR (Sigma-Aldrich, SML1046) was changed every two days. Number of organoids was counted on day 5, each experiment consisted in at least 4 technical replicates and was repeated at least two times.

### Tissue processing and Immunostaining

Intestinal jejunal segments were harvested, flushed with DPBS (Fisher Scientific, 14-190-250), cut longitudinally, and rolled around a cotton swab stick (Swissroll orientation). For cryo-preservation, tissue was fixed in 4% PFA (Electron Microscopy Sciences, 15714) overnight following 24h in 30% Sucrose (in PBS), and embedded in OCT. 5uM sections were stained with indicated Ab using standard experimental workflow. For immunohistochemistry tissue was fixed overnight in 10% buffered formalin (VWR, 89370-094) and embedded using standard experimental procedures. All embedding and sectioning was performed in the Molecular Pathology Shared Resource. IHC staining was performed using standard protocols. EdU was imaged using the Click-IT imaging assay (ThermoFisher, C10337).

### RNA extraction and qPCR

Single sorted cells (∼50’000 cells) were lysed in buffer RLT and stored at -80°C until starting the RNA isolation following manufacturer instructions (Qiagen, 74034). cDNA reaction was performed using qScript™ cDNA SuperMix (QuantaBio, 95048). qPCR analyses were performed using a QuantStudio 3 thermocycler (Applied Biosystems); gene expression is presented using the ΔΔCt method (*18S* or *Gapdh* genes used as housekeeping).

### 10x scRNAseq and single cell Multiome (ATAC+RNA)

Library preparation and sequencing were performed by the JP Sulzberger Columbia Genome Centre (Single cell analysis core), using standard methodologies. For scRNA-seq crypt epithelial cells were sorted to exclude dead cells and contaminating villi (Alive/Epcam+/B6A6-). Immediately after sorting, cells were counted using an automated cell counter (ThermoFisher Countess II FL) to check viability and processed for library preparation (10x chromium). For single cell Multiome, immediately after sorting, isolated cells (>100’000) were processed to extract nuclei following manufacturer recommendation (protocol CG000365, Rev B). Isolated nuclei were counted and quality control using an automated cell counter (ThermoFisher Countess II FL) and immediately processed for library preparation.

### Single-cell analysis of mouse intestinal crypt cells

Single-cell RNA-seq (scRNA-Seq) UMI profiles were processed using Seurat (v.4.1.0)^7^. Cells with >1,000 expressed genes and mitochondrial gene content < 10% were retained for downstream analysis, yielding to 3,656 cells. UMI counts were normalized and scaled using SCTtransform from the Seurat package^7^. Next, a Shared Neighbors Graph (SSN) was built with knn=10 to select cells with most similar transcriptional profiles and merge them to generate high resolution ensembles of cells called metacell: this approach augments the number of detected genes per cells, which usually is very low due to dropout technological bias (<20%), thus increasing the number of targets that can be recovered by reverse-engineering regulatory networks. Metacell profiles were computed on normalized data, but merged into UMI counts and transformed to count per million (cpm) for downstream analysis. Cell doublets were identified using scanpy’s implementation of scrublet^8^.

### Reverse-engineering of mouse intestinal crypt regulatory networks

An intestinal stem cell (ISC)-specific regulatory network (interactome) was reverse engineered from the resulting metacell cpm profiles (n = 1,218) using ARACNe-AP^9^, the most recent implementation of the ARACNe algorithm^10^, with 200 bootstraps, a Mutual Information (MI) P-value threshold P ≤ 10-8, and Data Processing Inequality (DPI) enabled. A total of n = 2305 regulatory proteins (RP) were selected into manually curated protein sets, including n = 1465 Transcription Factors (TF) and n = 840 co-Transcription Factors or chromatin remodeling enzymes, using the following Gene Ontology (GO) identifiers: GO:0003700 and GO:0003712^11, 12^. The resulting network includes 1,797 regulators, 14,935 targets and 548,442 interactions. Epithelial crypts scRNA-Seq profiles were transformed to protein activity profiles using the metacell-derived regulatory network and the VIPER algorithm ^13^. To avoid bias due to different regulon sizes, regulons were pruned to include only the 50 highest likelihood targets, as recommended in^13^, and regulons with < 50 targets were excluded from the analysis. Next, we sought to recover cell identities by identifying clusters of cells that share the same regulatory program using the Louvain clustering algorithm applied on the protein activity profiles of cells. Using these profiles, a SSN was built with knn=15 using the first 6 Principal Component (PC) as identified by the Elbow method. We performed a grid search analysis to tune Louvain’s resolution parameter to maximize the average of within-cluster Silhouette scores across each candidate optimal clustering solution. A high Silhouette score is an indication that clustered cells have homogenous profiles, hence as sampled from the same cell population. The optimal solution yielded 3 major clusters and differential markers analysis identified the two major intestinal lineages, secretory and absorptive, with the third population appearing to correspond to stem/progenitors, based on cell potency analysis (See Cell Potency Inference paragraph) For each one of the 3 clusters, a lineage-specific regulatory network was reverse-engineered as explained above and used to recover ISC cell identities.

### Lineage-specific protein activity analysis and identification of cell identities

Lineage-specific protein activity and clustering analysis was performed as follows. Crypt epithelial cells were computationally isolated in three distinct datasets based on their inferred lineage as identified at the previous step. For each lineage, a specific gene expression signature (GES) based on the relative subset of cells was computed by scaling the expression data after having normalized and variance-stabilized the UMI counts matrix using *sctransform* as implemented in the Seurat R package. Next, VIPER analysis was performed using cluster-specific regulatory networks and the lineage-specific GES as computed at the previous step. Clustering analysis was performed in a lineage-specific manner by re-running the grid search analysis on the isolated cells protein activity matrix. The absorptive lineage was divided in two clusters, the secretory lineage in three clusters and the stem/progenitor one in two clusters (see Extended Data). The cells from the whole crypt epithelial cell dataset were then labeled based on these seven clusters. To generate the final matrix of protein activity, a GES was generated by comparing each crypt epithelial cell to the average of all the cells independently of their lineage, and VIPER was run on each of the three lineage-specific regulatory networks for all the cell. For each cell, only the VIPER run with the best matching network was retained, as selected by enrichment of top 50 cell-specific candidate MRs. This ensures that each cell, especially those in transition, has been analyzed with its best matching lineage-specific network, and, that each cell is comparable with each other across the dataset because the reference distribution for the GES has been computed across all the crypt epithelial cells present in the dataset. For the RPs that were lineage specific, we used the VIPER score from the lineage-specific regulatory network analysis, while protein activity of RPs that were not present in one of the networks (e.g., due to their lineage specific activity) were computed using metaVIPER across all the networks ^14^.

### Display of crypt epithelial cells as scatter plot

For the purpose of displaying crypt epithelial cells on a 2-axis plane, we performed the following operations on the final VIPER matrix.

Specifically, a KNN graph of the data was built with k=15 using n=30 Principal Components (PCs). Next, we ran the partition-based graph abstraction (PAGA)^15^ algorithm on the data using the clustering solution we identified as input partition, to provide an informative initialization for the UMAP algorithm to increase its accuracy by initializing its embedding^16^. The UMAP embedding as computed with this protocol on protein activity data was then transferred to the gene expression matrix, along with information of clustering labels to generate the plots in the manuscript. Specifically, for DT-ablated and IR datasets, we performed asymmetric data integration with the *ingest* methodology included in the *scanpy* framework^17^, using CTRL dataset as reference.

### Analysis of publicly available intestinal epithelial crypt cells datasets

Dataset as UMI counts from *Ayyaz, et al.* (2019) and *Böttcher, et al.* (2021) where downloaded from GEO. For each dataset, GES was computed using an internal reference in each dataset independently, and metaVIPER was run using all the networks from the three lineages identified from the CTRL dataset. Cluster assignment on each cell was performed by enrichment analysis using top 50 candidate MR proteins for each of the 7 clusters. Specifically, a cell was assigned to the cluster for which it had the highest enrichment score.

### Cell Potency Inference

Cell potency inference was performed using three distinct approaches. The first one, the CytoTRACE algorithm, relies on gene expression data and leverages information embedded in the number of detected genes per cell^1^, to compute the CytoTrace Score (CT) that we used to sort cells from the least differentiated to the terminally differentiated ones. The second approach we used builds a one-class logistic regression (OCLR) using pluripotent stem cell bulk samples (ESC and iPSC) as a predictive model to compute a Stemness Index (SI) on new samples, that we used on protein activity profiles^2^. The third approach, SCENT, uses protein-protein interaction networks (PPIs) and single-cell transcriptomic data to infer signaling entropy scores (SR) to use as proxy of cell differentiation potency, with the hypothesis that a cell with high signaling entropy should be endowed with higher differentiation potential^18^. PPIs were modeled using PrePPI^19^, a large-scale database of human PPI, by retaining the top 5% of high confidence interactions. To convert murine gene products to human, for PrePPI network, gene identifiers were mapped to their human-mouse orthologs using R biomaRt services. To compute CT scores across pre- and post-radiation cells, we performed bootstrap analysis by subsampling 100 times the sham dataset with the same number of cells we recovered after irradiation, in order to make them comparable. The mean CT score per cell was used for the final analysis to compare cell potency in the post-radiation sample.

### Stem Cell Markers Discovery

Markers associated with high cell potency were inferred by correlating regulatory protein activity with CT scores. RP were prioritized based on the Pearson’s correlation coefficient p-value after correction for multiple hypothesis testing using the Benjamini-Hochberg method.

### Joint analysis of CTRL and IR-exposed crypt epithelial cells

To compare CTRL and IR datasets in a quantitative manner we build a null model of cell density estimates with a bootstrap approach and subsampling, to take also into account the difference of about an order of magnitude in sample size between CTRL (n∼3,500) and IR (n∼600). Specifically, differential Kernel Density Estimate (KDE) was performed in the following way. Pre- and post-radiation samples were normalized independently using sctransorm with method glmGamPoi. Next, the two datasets were joined together by identifying common anchors using reciprocal PCA as implemented in using Seurat (v.4.1.0) ^7^. metaVIPER analysis was performed using the three lineage-specific networks and the scaled integrated gene expression matrix as cell specific expression signature. Next, UMAP was performed on the protein activity matrix of the joined datasets using Euclidean as distance metric and 30 PCs. The first two UMAP dimensions were used to lay pre- and post-radiation cells on a bi-dimensional pane of 1e4 equally-sized tiles addressed by 100 intervals per UMAP dimension. The pre-radiation dataset was used to bootstrap 100 times a subset of 500 cells that were uniformly sampled to create a distribution of density estimates using a Gaussian kernel. To assess cell depletion or enrichment for each tile in the post-radiation sample, we used the positive or negative z-score computed using mean and standard deviation of the control (pre-radiation sample).

### Archetype analysis

Archetype analysis was performed in R using the *archetypes* package. Briefly, the VIPER matrix was analyzed using classical archetype analysis for every number of *k* archetypes between 2 and 12. The elbow method allowed to determine that k=5 is the optimal number of archetypes (see Extended Data).

### Paneth cells analysis

Paneth cells were identified in crypt epithelial cells by looking at the concurrent expression of all the following markers: *Lyz1*, *Mmp7*, *Dll4*, and *Nupr1,* calling it Paneth cell when all these markers were found expressed at any degree. This is a lower bound estimation of Paneth cells, given technical dropouts. Paneth cell specific candidate MR proteins were identified by selecting Paneth cells, generating a GES using as reference all the other non-Paneth cells, and running msVIPER using the secretory lineage regulatory network.

### scATAC-Seq analysis

scATAC-Seq data was processed using ArchR^20^. Peak calling was performed using MACS2 with default parameters^21^. Cluster analysis was performed using the RNA data modality by projecting the clustering solution identified on the CTRL sample using *ingest* over scanpy^17^. Known motifs enrichment analysis was performed using the *cisbp* motifs dataset and the 7-cluster solution identified over the VIPER-transformed data.

### Data availability

scRNAseq datasets together with single cell Multiome (scRNA+scATAC seq) of crypt epithelial cells will be made available before publishing these results.

### Code availability

All code will be available on Github or upon request at the moment of publication.

